# Life on the leaf: Seasonal activities of the phyllosphere microbiome of perennial crops

**DOI:** 10.1101/2021.04.20.440608

**Authors:** Adina C. Howe, Nejc Stopnisek, Shane K. Dooley, Fan Yang, Keara L. Grady, Ashley Shade

## Abstract

Plants and microorganisms form beneficial associations. Understanding plant-microbe interactions will inform microbiome management to enhance crop productivity and resilience to stress. Here, we apply a genome-centric approach to identify key leaf microbiome members on field-grown switchgrass and miscanthus, and quantify their activities for switchgrass over two growing seasons. We integrate metagenome and metatranscriptome sequencing from 192 leaf samples collected over key time points in crop phenology. We curated 40 focal metagenome-assembled-genomes (MAGs) and conservatively focus analysis on transcript recruitment to medium and high-quality MAGs that were <10% contaminated and >50% complete. Classes represented by these MAGs (Actinomycetia, Alpha- and Gamma-Proteobacteria, and Bacteroidota) were active and had seasonal dynamics in key functions, including enrichments in transcripts for of short chain dehydrogenase, molybdopterin oxioreductase, and polyketide cyclase in the late season. The majority of MAGs had activated stress-associated pathways, including trehalose metabolism, indole acetic acid degradation, betaine biosynthesis, and reactive oxygen species degradation, suggesting direct engagement with the host environment. We also detected seasonally activated biosynthetic pathways for terpenes (carotenoid and isoprenoids), and for various non-ribosomal peptide pathways that were poorly annotated. Overall, this study overcame laboratory and bioinformatic challenges associated with field-based leaf metatranscriptome analysis to inform potential key activities of these phyllosphere populations. These activities collectively support that leaf-associated bacterial populations are seasonally dynamic, responsive to host cues and interactively engage in feedbacks with the plant.

## Introduction

Perennial plants are a crucial target for the sustainable development of biofuels (Robertson *et al*., 2017; Hestrin *et al*., 2021; Ma *et al*., 2021). In addition to yielding high biomass that can be converted to biofuels and bioproducts, perennial crops offer a broad range of ecosystem services that support efforts to mediate climate change, including greenhouse gas mitigation and promotion of soil nutrient cycling (Heaton *et al*., 2008; Langholtz *et al*., 2016; Robertson *et al*., 2017; Roley *et al*., 2018). Like all plants, perennials harbor diverse microbiota, and many of these microbes are either known or expected to benefit their hosts. For example, plant-associated microbes can increase productivity and protect against environmental stressors. Because of the intimate engagement of many plant-associated microbiota with the host, management of the plant microbiome is one tool proposed to promote crop vigor and support crop resilience to global climate changes (Busby *et al*., 2017; Toju *et al*., 2018; Haskett *et al*., 2020; Wang and Haney, 2020). Therefore, along with selective breeding and data-informed field management, regulating the plant microbiome is expected to be key for the sustainable production of perennial biofuel feedstocks.

Plants have anatomical compartments that each are inhabited by distinctive microbial consortia. Generally, the diversity and composition of the plant microbiome narrows from external compartments to internal, and the plant plays an active role in filtering the microbiome composition inward (Hacquard *et al*., 2015; Hardoim *et al*., 2015; Gopal and Gupta, 2016). External plant compartments include the root zone, rhizosphere and rhizoplane below ground, and the epiphytic phyllosphere above ground (Andrews and Harris, 2000). External compartments have relatively higher representation of transient or commensal microbial taxa, and these compartments engage with and recruit microbes from the immediate environment. Internal compartments include the endosphere of above- and below-ground tissues, and these have relatively low richness and harbor the most selected microbiota (Bulgarelli *et al*., 2013; Müller *et al*., 2016). Of these compartments, the rhizosphere has received the most attention as a key site of microbial-plant interactions that are important for nutrient and water acquisition (e.g., Kuzyakov and Razavi, 2019). However, members of the microbiota that inhabit the phyllosphere also provide important plant functions, such as pathogen exclusion and immune priming (Bell *et al*., 2019; Chen *et al*., 2020). Phyllosphere microorganisms have specialized adaptations to their exposed lifestyle (Lindow and Brandl, 2003; Vorholt, 2012; Müller *et al*., 2016; Koskella, 2020) and contribute to global carbon and other biogeochemical cycling, including transformations relevant for climate change (Bringel and Couée, 2015; Dorokhov *et al*., 2018; Cavicchioli *et al*., 2019). Because perennial biofuel feedstocks are selected to maximize foliage surface area, understanding the phyllosphere microbiome is expected to provide insights into key microbial engagements that benefit the plant to support productivity and stress resilience.

There are two general challenges in regulating the microbiome to promote crop vigor and resilience to environmental stress. The first challenge is to distinguish the beneficial members of the plant microbiome from transient or commensal members, with recognition that some members that provide host benefit likely change situationally, either over plant development or given environmental stress (Edwards *et al*., 2015; Xu *et al*., 2018; Zhalnina *et al*., 2018), while others are stable (Shade and Stopnisek, 2019; Stopnisek and Shade, 2021). The second challenge is that plants and their agroecosystems are temporally dynamic over the growing season, and their associated microbiota are also dynamic. It is currently unclear what functions may be associated with phyllosphere microbial dynamics and their potential interactions with plant hosts.

Previously, we used 16S rRNA gene amplicon analysis to identify a “core” cohort of bacterial and archaeal taxa that were persistently associated with the phyllosphere microbiomes’ of two perennial biofuel feedstocks, miscanthus and switchgrass. Persistent membership was established by collecting leaf samples over replicated field plots, over a temperate seasonal cycle, and across two annual growing seasons for switchgrass (Grady *et al*., 2019). Other studies in switchgrass have similarly reported that the leaf, and other plant compartments, can be distinguished by microbiome compositions (e.g., Bulgarelli *et al*., 2012; Lundberg *et al*., 2012; Bahulikar *et al*., 2014; Roley *et al*., 2019; Singer *et al*., 2019), suggesting selection to the leaf compartment. In the current study, we aimed to understand the functional attributes and activities of persistent phyllosphere taxa, with an interest in specialized adaptations to the leaf and interactions with the host plant that may inform the mechanisms and nature of plant-microbe engagements. Therefore, we performed a seasonal analysis of phyllosphere metagenomes for both miscanthus and switchgrass. We paired our metagenome longitudinal series with metatranscriptome analyses at select time points in switchgrass phenology to determine which functions were active seasonally. We performed genome-centric analyses of the metagenome and focused on understanding the seasonal dynamics and functions of a focal subset of medium- and high-quality metagenome-assembled-genomes (MAGs) that we could bin from these data. Our results reveal functions supportive of a leaf-associated lifestyle and seasonal activities of persistent phyllosphere members. Finally, we provide evidence that these genomes were detected in various sites and years beyond our original study plots, suggesting that they are general, consistent inhabitants of bioenergy grasses.

## Materials and Methods

### Site description and sampling scheme

Switchgrass and miscanthus leaves and corresponding contextual data were collected at the Great Lakes Bioenergy Research Center (GLBRC) located at the Kellogg Biological Station in Hickory Corners, MI, USA (42°23’41.6” N, 85°22’23.1” W) (**Figure 1**). We sampled switchgrass (*Panicum virgatum L.* cultivar Cave-in-rock) and miscanthus (*Miscanthus x giganteus*), from the Biofuel Cropping System Experiment (BCSE) sites, plot replicates 1-4, as previously described (Grady *et al*., 2019). This included collecting leaves from switchgrass and miscanthus at 8 and 9 time points, respectively, in 2016 and switchgrass at 7 time points in 2017 seasons (**Table 1, Figure 1, Dataset 1, Dataset 2**). We collected leaves for RNA isolation at three phenology-informed switchgrass time points in 2016 (emergence, peak growth and senescence according to GLBRC standard phenology methods https://data.sustainability.glbrc.org/protocols/165) to assess the potential for sufficient mass and quality RNA extraction from the switchgrass leaf surface, and then expanded to include leaves from all switchgrass sampling time points in 2017. Leaves for RNA isolation were flash-frozen in liquid nitrogen immediately and stored at −80°C until processing.

**Figure 1.**
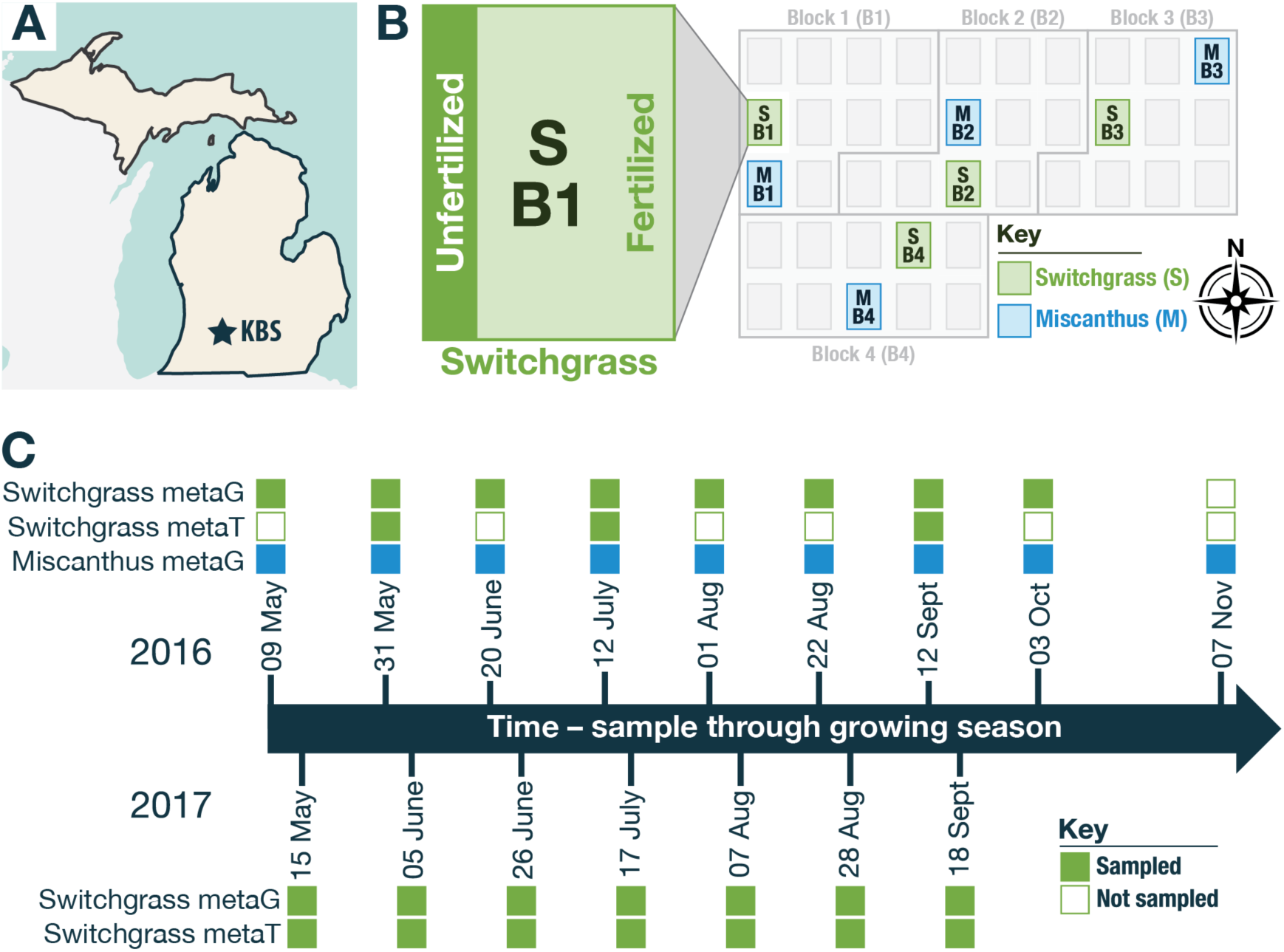
Phyllosphere microbiome field sampling strategy at the Great Lakes Bioenergy Research Center Bioenergy Cropping System Experiment (BCSE) in 2016 and 2017. (A) The study site is at Kellogg Biological Station, a Long-Term Ecological Research site focused on agroecosystems located in southwest Michigan. (B) Four replicate randomized cropping system blocks from the BCSE were sampled at each time point for switchgrass and/or miscanthus, and within each plot there was a fertilized main plot and unfertilized subplot sampled. (C) In 2016, both switchgrass and miscanthus were sampled, and in 2017 only switchgrass was sampled. Switchgrass leaves were flash frozen in liquid nitrogen for RNA extraction and metatranscriptome analysis at a subset of time points in 2016 and at all points in 2017.

**Table 1.** Summary of RNA and DNA samples that returned reads and passed Illumina standard quality control at the Joint Genome Institute. Total collected samples submitted for sequencing is provided first, and the number of quality sequencing datasets returned for analysis is given in parentheses.

|  | 2016 |  | 2017 |  |
| --- | --- | --- | --- | --- |
|  | Metagenome | Metatranscriptome | Metagenome | Metatranscriptome |
| Switchgrass | 8 timepoints<br>(64/64<br>successful) | 3 (22/24<br>successful) | 7 (56/56<br>successful) | 7 (56/56<br>successful) |
| Miscanthus | 9 (72/72<br>successful) | Not assessed | Not assessed | Not assessed |

### Phyllosphere RNA and DNA isolation

Phyllosphere epiphyte DNA was isolated and processed (Grady *et al*., 2019). DNA concentrations were normalized to 4 ng/ml. Phyllosphere epiphyte RNA was isolated using a benzyl chloride liquid:liquid extraction, based on (Grady *et al*., 2019) that was newly modified for RNA isolation based on the published methods of (Suzuki *et al*., 2001). Approximately 5 g of intact, frozen leaf material was added to a 50 ml polypropylene conical tube (Corning #430290) and kept frozen on liquid nitrogen while samples were weighed and transferred. Denaturing Solution (DS) was prepared with 4.2 M guanidine thiocyanate, 25 mM sodium citrate dihydrate pH 7.0, 0.5% (v/v) sodium n-laroyl sarcosine in Milli-q water and was filter sterilized through 0.22 mm filters. Immediately prior to extraction, a working stock of DS was prepared fresh by adding 2-mercaptoethanol to a final concentration of 5% (v/v, DS/2-ME). To each leaf tube, 5 ml of benzyl chloride, 2.5 ml of 3M sodium acetate (pH 5.2), and 5 ml of the working stock of DS/2-ME was added. The tube was incubated in a 60°C water bath for 20 minutes with vortexing every 1 min. The leaves were removed from the conical tube using ethanol-sterilized forceps and discarded.

Five ml of chloroform:isoamyl alcohol (24:1) were added to the remaining solution in each conical tube, which were then shaken by hand for 15 seconds and incubated on ice for 15 minutes. The tubes were then centrifuged at 12,000 x g for 15 min at 4°C to separate aqueous and organic phases. Up to 5 ml of the upper, aqueous phase was transferred to a clean 15 ml polypropylene tube, without disrupting the white interface. Two and a half ml of sodium citrate buffer (1.2 M sodium chloride, 0.8 M sodium citrate dihydrate in Milli-q water, filter sterilized at 0.22 mm) and ice-cold isopropanol were added to achieve a final volume of 12.5 ml. Next, the tubes were centrifuged at 12,000 x g for 15 minutes to pellet the RNA, and the remaining supernatant aspirated using a pipette. The RNA pellets were resuspended in 0.3 ml of working DS/2-ME solution, and afterwards 0.3 ml of ice-cold isopropanol was added and mixed by pipetting gently. The solution was incubated for 30 minutes at −20°C, and then the full volume was transferred to a clean nuclease-free 1.7 ml tube and centrifuged at 16,000 x g for 15 minutes at 4 °C. The supernatant was removed, and the pellet washed in 1 ml of nuclease-free 75% ethanol. Tubes were then centrifuged at 16,000 x g for 15 minutes at 4 °C, and supernatant again removed using a pipette. The remaining pellet was air dried to completely remove residual ethanol, then resuspended in 30 ml of nuclease-free Tris-EDTA buffer, pH 8.0.

Genomic DNA (gDNA) was removed using RNase-free DNase I (Thermo Fisher #AM2222) per manufacturer’s instructions. The RNA was then purified using the RNeasy MinElute Cleanup Kit (Qiagen Germantown, MD, USA) according to manufacturer’s instructions. The absence of contaminating gDNA was confirmed by lack of amplification of the 16S rRNA gene V4 region by PCR (Caporaso *et al*., 2011) with positive and negative controls. This RNA isolation method was developed to most closely align with our established phyllosphere epiphyte DNA isolation (Grady *et al*., 2019) in order to minimize potential bias introduced during biofilm disruption or microbial cell lysis, as well as to minimize contamination from host RNA or genomic DNA by leaving the plant tissue intact. Commercial RNA extraction kits are primarily based on grinding or bead beating whole tissue samples, which would result in over-representation of host-derived nucleic acids and potentially introduce bias in microbial cell lysis efficiencies.

### Metagenome and metatrascriptome library preparation

The Joint Genome Institute (JGI) performed the library preparation and sequencing from submitted RNA and DNA samples. Plate-based DNA library preparation for Illumina sequencing was performed on the PerkinElmer Sciclone NGS robotic liquid handling system using Kapa Biosystems library preparation kit. 1.82 ng of sample DNA was sheared to 436 bp using a Covaris LE220 focused-ultrasonicator. The sheared DNA fragments were size selected by double-SPRI and then the selected fragments were end-repaired, A-tailed, and ligated with Illumina compatible sequencing adaptors from IDT containing a unique molecular index barcode for each sample library. The prepared libraries were quantified using KAPA Biosystems’ next-generation sequencing library qPCR kit and run on a Roche LightCycler 480 real-time PCR instrument. Sequencing of the flowcell was performed on the Illumina HiSeq sequencer following a 2×151 indexed run recipe.

At JGI, plate-based RNA sample prep was performed on the PerkinElmer Sciclone NGS robotic liquid handling system using Illumina Ribo-Zero rRNA Removal Kit (Bacteria) and the TruSeq Stranded Total RNA HT sample prep kit following the protocol outlined by Illumina in their user guide: https://support.illumina.com/sequencing/sequencing_kits/truseqstranded-total-rna.html, and with the following conditions: total RNA starting material of 100 ng per sample and 10 cycles of PCR for library amplification. The prepared libraries were quantified using KAPA Biosystems’ next-generation sequencing library qPCR kit and run on a Roche LightCycler 480 real-time PCR instrument. Sequencing of the flowcell was performed on the Illumina NovaSeq sequencer using NovaSeq XP V1 reagent kits, S4 flow cell, following a 2×151 indexed run recipe.

### Quality filtering of metagenomes and metatranscriptomes

We proceeded with bioinformatic analysis (**Figure 2**) of 192 metagenome (**Dataset 1**) and 78 metranscriptome (**Dataset 2**) observations that met JGI standards for raw data quality based on the Illumina proprietary software. We used Trimmomatic (v0.39) (Bolger *et al*., 2014) to remove adaptors and filter low-quality reads from fastq files using the following arguments: PE-phred33 ILLUMINACLIP:TruSeq3-PE-2.fa:2:30:10:8:TRUE LEADING:3 TRAILING:3 SLIDINGWINDOW:4:15 MINLEN:36. After assembly, plant host reads were filtered out (for both metagenomes and metatranscriptomes) by removing all reads that mapped to the switchgrass genome (*Panicum virgatum* v1.0, DOE-JGI, http://phytozome.jgi.doe.gov/) or miscanthus genome (*Miscanthus sinensis* V7.1 http://phytozome.jgi.doe.gov/) using bowtie2 (v 2.4.1), samtools (v 1.13), and bedtools (v2.30.0) (Li *et al*., 2009; Quinlan and Hall, 2010; Langmead and Salzberg, 2012) (**Figure S1)**. To remove the fungal reads and improve the prokaryotic signal in metagenome samples, we also filtered reads against 7 fungal genomes (Gostinčar *et al*., 2014; Druzhinina *et al*., 2018; Gill *et al*., 2019; Haridas *et al*., 2020) that represent close relatives of the most abundant fungal species in this system that we assessed and reported in our prior work (Bowsher *et al*., 2020) (**Table S1**). The genomes of these fungal species were retrieved from the JGI Genome Portal.

**Figure 2.**
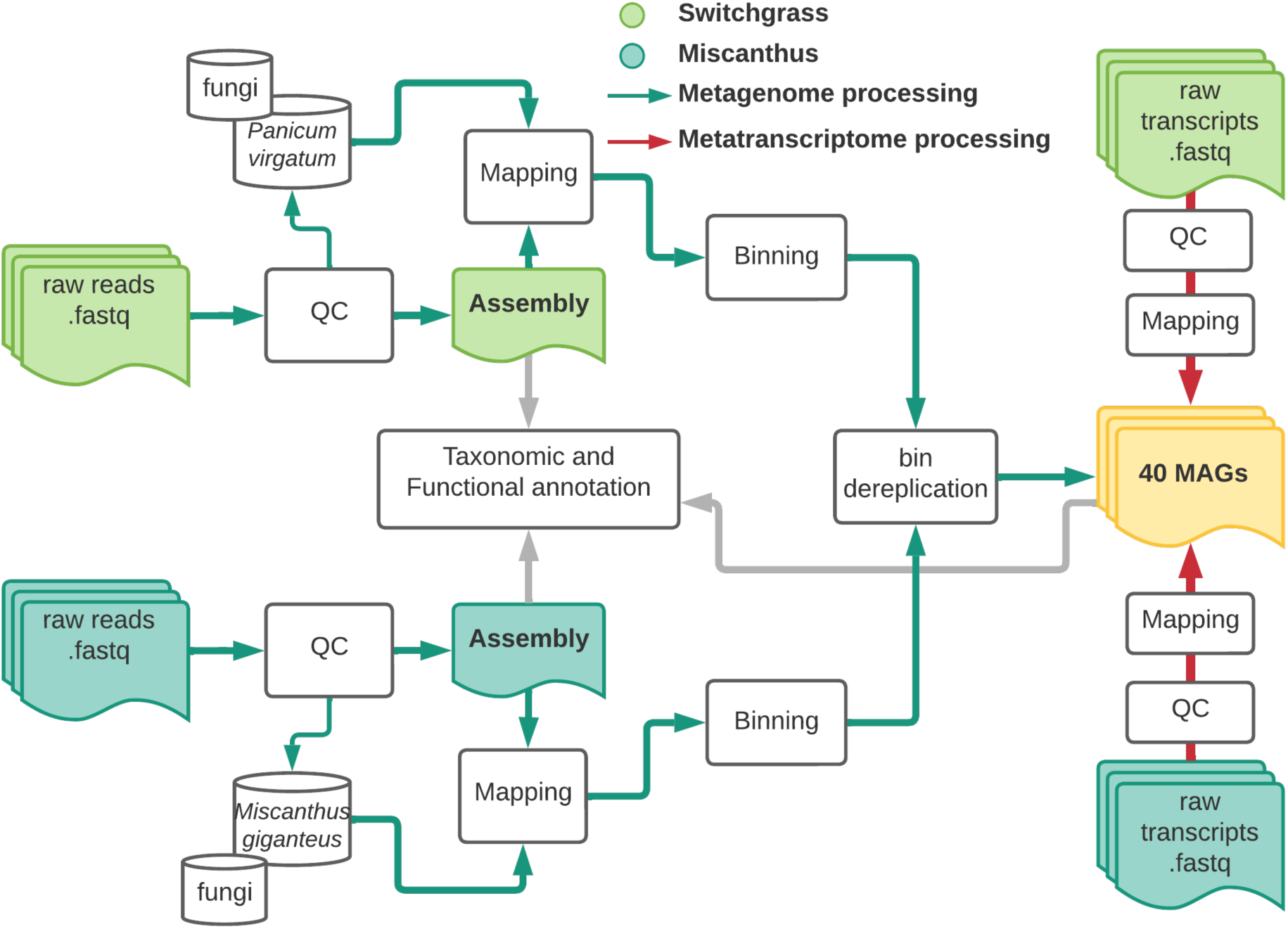
Overview of bioinformatic processing of the metagenome (green solid arrows) and metatranscriptome (red solid arrows) datasets. Switchgrass reads are shown in light green and miscanthus is dark green. The solid grey arrow represents the common analysis step for both datasets. Figure was made with Lucidchart.

### Metagenome assemblies and metagenome-assembled genome binning, curation, refinement, annotation

Two metagenome assemblies, one for switchgrass and one for miscanthus, were created based on metagenomes collected in 2016. These filtered metagenome reads were combined and used for co-assembly with MEGAHIT (v 1.2.9) using--kmin-1pass (low sequencing depth) and--presets meta-large (complex metagenome) (Li *et al*., 2015). Additionally, we curated metagenome assembled genomes (MAGs) from the 2016 switchgrass and miscanthus metagenome libraries (n = 136 metagenomes) using Metabat (v2.2.15) (Kang *et al*., 2019). MAG assemblies were performed using filtered reads from switchgrass and miscanthus sampled from 2016, separately, to maximize completeness and reliability, as also done in other studies (Nayfach *et al*., 2019). To assess the MAGs quality and completeness, we used CheckM (v.1.13 with the lineage_wf option) estimates of quality and completeness (Parks *et al*., 2015). Among 238 MAGs assembled from the switchgrass or miscanthus phyllosphere metagenomes (**Dataset 3**), we selected a subset of MAGs based on: completeness greater than 50% and contamination less than 10%. We identified replicated bins associated with MAGs using dRep (v3.2.0,Olm et al. 2017), resulting in the removal of a single MAG. An additional MAG was removed due to insufficient read recruitment in metatranscriptomes (described below), for a final total of 40 focal MAGs, including 7 high-quality and 33 medium quality (Bowers *et al*., 2017) (**Dataset 3).**

Read recruitment was performed with MAGs that meet either medium- and/or high-quality standards (Swan *et al*., 2013; Bowers *et al*., 2017). The metagenome abundance of contigs in MAGs in each sample was estimated based on the median coverage of filtered metagenome reads associated to each MAG contig. Specifically, Bowtie2 (v2.2.2) was used to align reads to all focal MAG contigs (using default setting and allowing for a single read to map to only one reference). Bedtools (v2.28) was used to estimate the coverage of each basepair within the contig. The estimated abundance of contig was based on the median basepair coverage of all reads mapped to the contig, and the estimated abundance of a MAG was based on the average median coverage of all contig within its bins. The metatranscriptome abundance was estimated based on protein-encoding genes identified in MAGs. For each MAG contig, open reading frames (ORFs) and functional genes were identified using Prodigal (v2.6.3, default parameters). Transcripts were mapped to ORFs associated with each MAG to estimate median base coverage of each ORF (Bowtie2, default parameters, no multiple mappings allowed). To normalize varying sequencing depths, estimated abundances were normalized by the sum of the median base pair coverage of housekeeping genes identified in each sample. Housekeeping genes were identified based on full sequence alignment to the HMM models of 71 housekeeping single-copy genes with E-value of less than 1e-5 (Eren *et al*., 2015; Lee, 2019). If housekeeping genes were not identified in a metagenome or metatranscriptome, samples were removed from further analysis. We also estimated the total reads associated with each MAG for metagenomes and metatranscriptomes (**Dataset 4**). One MAG, M22, had an average of 48 reads map to metatranscriptome and was removed for further analysis. For metatranscriptome analyses, only ORFs with the top 75% of observed abundances and in at least 10% of samples were considered.

Functional annotation of ORFs in focal MAGs was done with the DRAM tool (v1.1.1, (Shaffer *et al*., 2020) using UniRef90, MEROPS, PFAM, dbCAN-HMMdb (v8) databases (all compiled with DRAM on February 12, 2021), and the KEGG database which we manually added to the DRAM pipeline (release January 1, 2018). To obtain functions related to terpene metabolism, ORFs associated with any KEGG annotation that contained the phrase “terpen” were selected.

MAGs were assigned taxonomy using GTDB-tk (v1.4.0, (Parks *et al*., 2018)). Assembled focal MAGs were aligned againts the chloroplast genome of *Panicum virgatum* (NC015990) (BLAST, 2.10.1). We detected partial matches to eight bins and no full alignments, confirming that those bins were not chloroplast genomes. Additionally, we compared the focal MAGs taxonomy to our previously detected 16S rRNA gene core cohort (Grady *et al*., 2019). This cohort consists of 61 phylogenetically diverse bacterial OTUs (97% clustering). Taxonomy of these 61 OTUs was compared with the 40 MAGs at the genus level only. Due to differences in nomenclature between the GTDB and SILVA databases we removed extensions for MAGs classified as *Pseudomonas_E* or *Aeromonas_A*.

### Statistical analyses

Pairwise comparisons of functional roles (the cumulative sum of all associated ORFs) between the early (May – June) and late (July – Sep) season were performed with the Kruskal-Wallis test by ranks. Early and late season were delineated by plant phenology (flowering/senescence in late season). The distribution and dynamics of MAGs’ transcripts were compared with Ward’s method for hierchical clustering using euclidian distances of the estimated metatranscriptome abundances using *pvclust* (version 2.20)(Suzuki and Shimodaira, 2006).

### Assessment of biosynthetic gene clusters (BGC) on focal MAGs

Biosynthetic gene clusters (BGC) were predicted by antiSMASH (v6.0) (Blin *et al*., 2021) and further annotated by Big-SCAPE (v1.1.0) (Navarro-Muñoz *et al*., 2020). While there are only 8 BGC classes used in Big-SCAPE (i.e. PKS I, PKS other, NRPS, RiPPs, Saccharides, Terpenes, PKS/NRPS Hybrids, Others), antiSMASH provides more detailed classification. We leveraged both outputs to investigate the diversity and expression of predicted BGCs in focal MAGs. From each predicted gene cluster, we extracted the location of the biosynthetic gene (i.e., gene_kind=”biosynthetic” and core_position). To evaluate the transcription of putative BGCs within a MAG, we searched for any transcripts that are mapped to the same genomic region of predicted biosynthetic genes.

### Predicting functional potential of focal MAGs

We used gapseq (Zimmermann *et al*., 2021) to predict completed metabolic pathways in our focal MAGs. We used the ‘find -p all –b 200’ option to search for pathways against the MetaCyc database. We filtered out incomplete pathways and remaining pathways were grouped into broader categories using MetaCyc classification and manually curated to focus on understanding the pathways relevant either to the plant environment or for microbial interactions with plants. These categories were defined as potential involvement in: i) plant (using plant metabolites/cell components), ii) phytohormone (known/potential involvement in phytohormone homeostasis), iii) stress (e.g., drought, reactive oxygen species), and iv) general (pathways that utilize potential plant derived products). Furthermore, we also manually searched for genes/pathways that were missed by gapseq but are known to be related to adaptation to plant-associated lifestyle, including secretion systems (Tseng *et al*., 2009; Lucke *et al*., 2020), oxidation of trace gases (H_2_ and CO)(Bay *et al*., 2021; Palmer *et al*., 2021), oxalate degradation, and phytohormone production/degradation. Similar to our biosynthetic gene analysis, the activity of predicted pathways was estimated based on mapped transcripts to key identified genes (**Table S2**). Contigs of focal MAGS were aligned (BLAST v2.7.1+) against nine isoprenoid precursor biosynthesis genes previously reported (Julsing *et al*., 2007). The top hit of each query was considered and accepted as aligned if E-value was less than 1e-5.

### Detection of MAGS in public metagenomes

To evaluate the presence of MAGs in other switchgrass and miscanthus metagenomes, 55 publicly available switchgrass, miscanthus, and corn metagenomes were used (**Dataset 5**). Metagenome reads were mapped to focal MAGs, using the same methods for mapping and abundance estimation described above for the metagenomes generated in this study.

### Data and code availability

Metagenome and metatranscriptome data are available through the Joint Genome Institute Genome Portal under proposal ID 503249. Code and links to data and metadata are available on GitHub (https://github.com/ShadeLab/PAPER_Howe_2021_switchgrass_MetaT). MAGs are available in NCBI under project IMG. All metadata, including metadata standards for metagenomes, metatranscriptomes, and metagenome-assembled genomes are provided as supporting datasets (**Datasets 1-6**).

## Results

### Key leaf bacterial populations and general dynamics

Among all assembled MAGs (n=238), we focused analysis on 40 that were high- and medium-quality based on completeness and contamination standards (**Figure 3A**, **Dataset 3**)(Bowers *et al*., 2017). The consistent detection of these 40 MAGs in both miscanthus and switchgrass samples (**Figure 4**) suggests that their originating populations were not host-specific but rather distributed among these perennial grasses; this supports our previous results from an 16S rRNA gene amplicon survey that revealed substantial overlap in the major leaf bacterial taxa across the two crops (Grady *et al*., 2019). Furthermore, there were 38 “core” OTUs from our prior survey that were also found in 25 focal MAGs here per their taxonomic identifications (**Figure 3B**); our previous study prioritized OTUs based on abundance and seasonal and field-replicated occupancy, and so this conservatively shows that at least half of the focal MAGs represent the most abundant and consistently detected taxa in this ecosystem. The 40 focal MAGs most represented the orders Rhizobiales (n=8/40), Actinomycetales (n=6/40), Burkholderiales (n=6/40), and Sphingomonadales (n=6/40) (**Dataset 3**).

**Figure 3.**
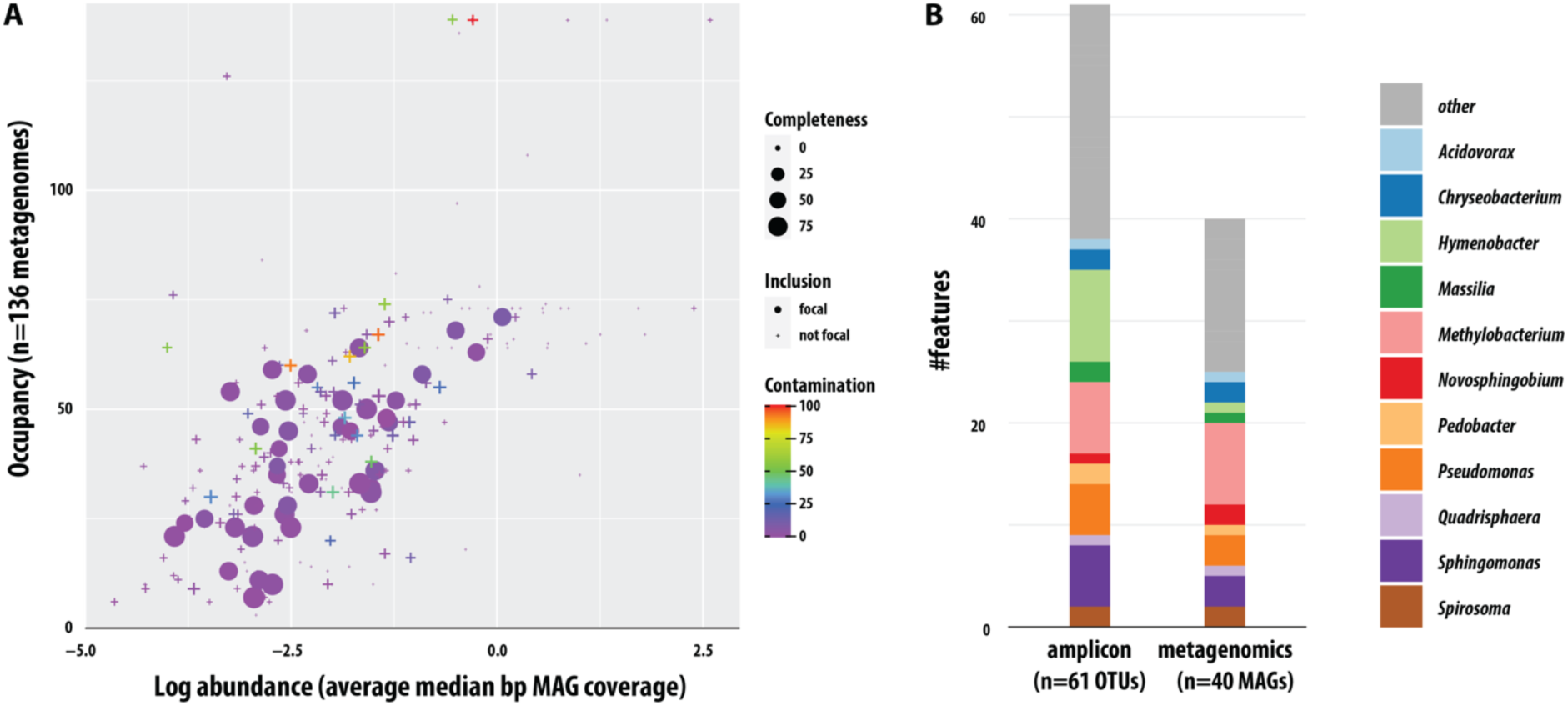
Summary of leaf-associated MAGs. **(A)** Abundance and occupancy of genomes assembled and binned from switchgrass and miscanthus phyllosphere metagenomes. Quality and contamination assessment were determined using checkM. 40 focal MAGs were selected **(B)** Taxonomy of focal MAGs, as annotated with GTDB-tk, and taxonomic overlap with the persistent taxa detected in our previous 16S rRNA gene amplicon survey.

**Figure 4.**
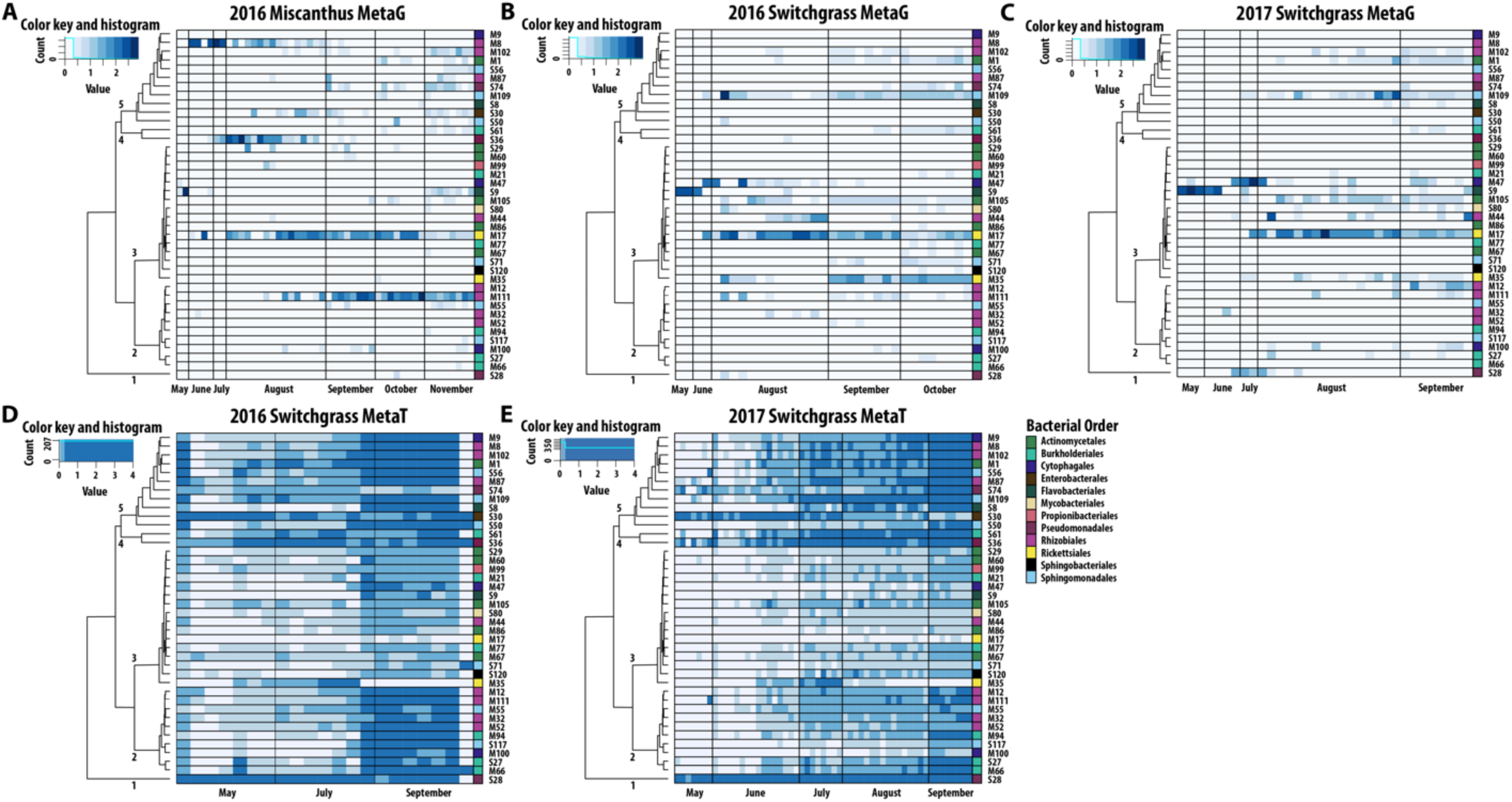
Seasonal patterns of the 40 focal MAG metagenome and metatranscriptome read recruitment. MAG abundances for: (**A**) Miscanthus 2016 metagenomes (metaG); (**B**) Switchgrass 2016 metagenomes; (**C**) Switchgrass 2017 metagenomes; (**D**) Switchgrass 2016 metatranscriptomes (metaT) and (E) Switchgrass 2017 metatranscriptomes. Abundances of metagenome contigs were estimated with the median basepair of read recruitment divided by the average median basepair coverage of housekeeping genes. Abundances of metatranscriptome ORFs were estimated based on the median basepair coverage of all reads mapped to ORFs and divided by median basepair coverage of housekeeping genes. The same dendrogram is applied to all panel, and it is the result of hierarchical clustering (see Figure S2) of metatranscriptome diversity and abundances in switchgrass.

Focal MAGs exhibited varied seasonal patterns in their overall metagenome (**Figure 4A-C**) and metatranscriptome (**Figure 4D-E**) read recruitment, with early, late, and consistent seasonal detection observed. However, in the metatranscriptomes of both years for switchgrass, there was a strong seasonal enrichment over time, with increases in the later months of sampling. Phenologically, August samples corresponded with (late) peak biomass and fruiting for switchgrass, and with a closed canopy for both switchgrass and miscanthus. Senescence occurred as early as mid-September for switchgrass and as late as November for miscanthus. Notably, transcript recruitment to the MAGs was robust even when there was inconsistent detection in metagenome read recruitment, suggesting that these MAGs were consistently active and had relatively high activity even when their genome detection was obscured by the flanking community or host signal.

MAGs were clustered based on coherence in seasonal transcript dynamics (**Figure 4D-E, Figure S2**). Five coherent and statistically supported groups (clusters) of 40 MAGs were identified (**Figure S2**). Cluster 1 contained a single MAG S28, assigned to genus *Pseudomonas* (S28, 98% complete, 1.9% contamination) and its genes were enriched early in the season (2017) but the MAG was highly active throughout the season. The other four clusters contained numerous MAGs and were relatively more dynamic, with trends towards late-season enrichment in transcripts. Several clusters contained MAGs annotated to the same class, suggestive of taxonomic coherence in seasonal activities. For example, cluster 3 included all but one (of eight) Actinomycetia MAGs.

### MAG genes and activities support a leaf-associated lifestyle

Not surprisingly, the most abundant subsystems identified among the 40 focal MAGs were associated with bacterial growth, such as carbohydrates, energy, amino acid, and nucleotide metabolisms (**Figure 5**). Leaf metatranscriptomes also showed that most subsystem transcripts were steadily enriched over the season. However, subsystem associated with colonization and adaptation, such as cell motility, signal transduction, community-associated pathways, and environmental adaptation trended down in their normalized transcripts over the season. A closer investigation of these down-trending functions revealed that they could be attributed almost exclusively to MAG S28 (*Pseudomodales* from cluster 1 in **Figure 4DE** and **Figure S3**). Finally, a few KEGG classifications were relatively stable and lacked overall seasonal trends, including transcripts for translation, transcription, and antimicrobial resistance.

**Figure 5.**
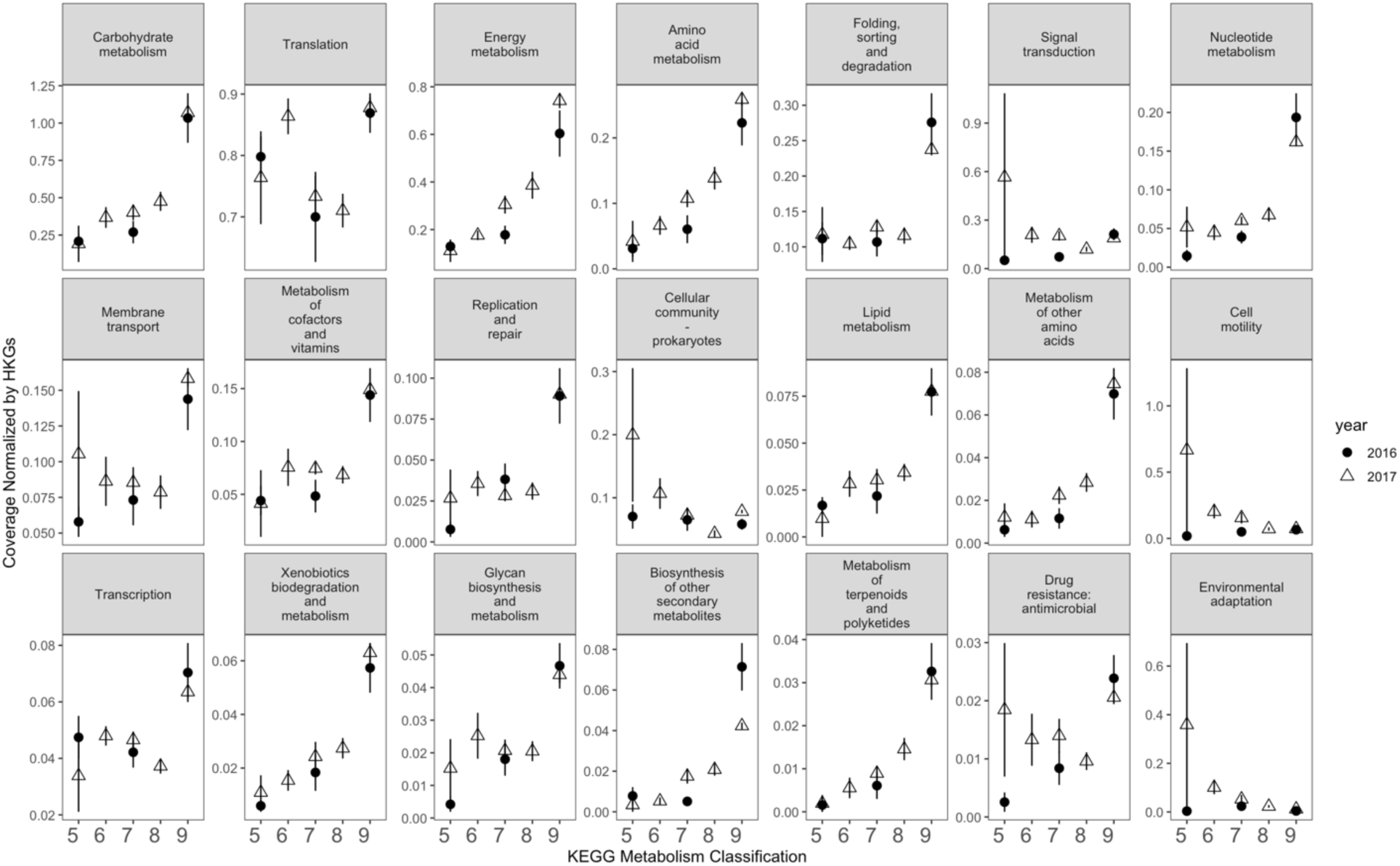
2016 (circle) and 2017 (triangle) switchgrass leaf transcript dynamics of KEGG metabolism classifications to the 40 focal MAGs. The y-axis is scaled for each classification.

We selected the most prominent phyllosphere transcripts based on their high detection among the 40 focal MAGs, hypothesizing that some of these functions shared between bacterial populations may represent functions for survival on the leaf. A total of 124 distinct functional roles were identified in at least 39 of the 40 MAGs (**Dataset 6, Figure S4**). Of these, most functions were associated to translation (n=48/124). Other broadly present functions were associated with metabolism of energy (n=29/124) and carbohydrates (n=28/124). There were also several functional roles associated with cofactors and vitamins (n=18/124). While many functions were enriched in transcripts in the late season (July-Sept) relative to the early season (May-June), three functional roles in particular stood out, with enrichments greater than 20-fold in the late season transcripts and included proteins classified as short chain dehydrogenase, molybdopterin oxidoreductase, and polyketide cyclase (**Table 2**, **Dataset 6, Figure S4**).

**Table 2.**
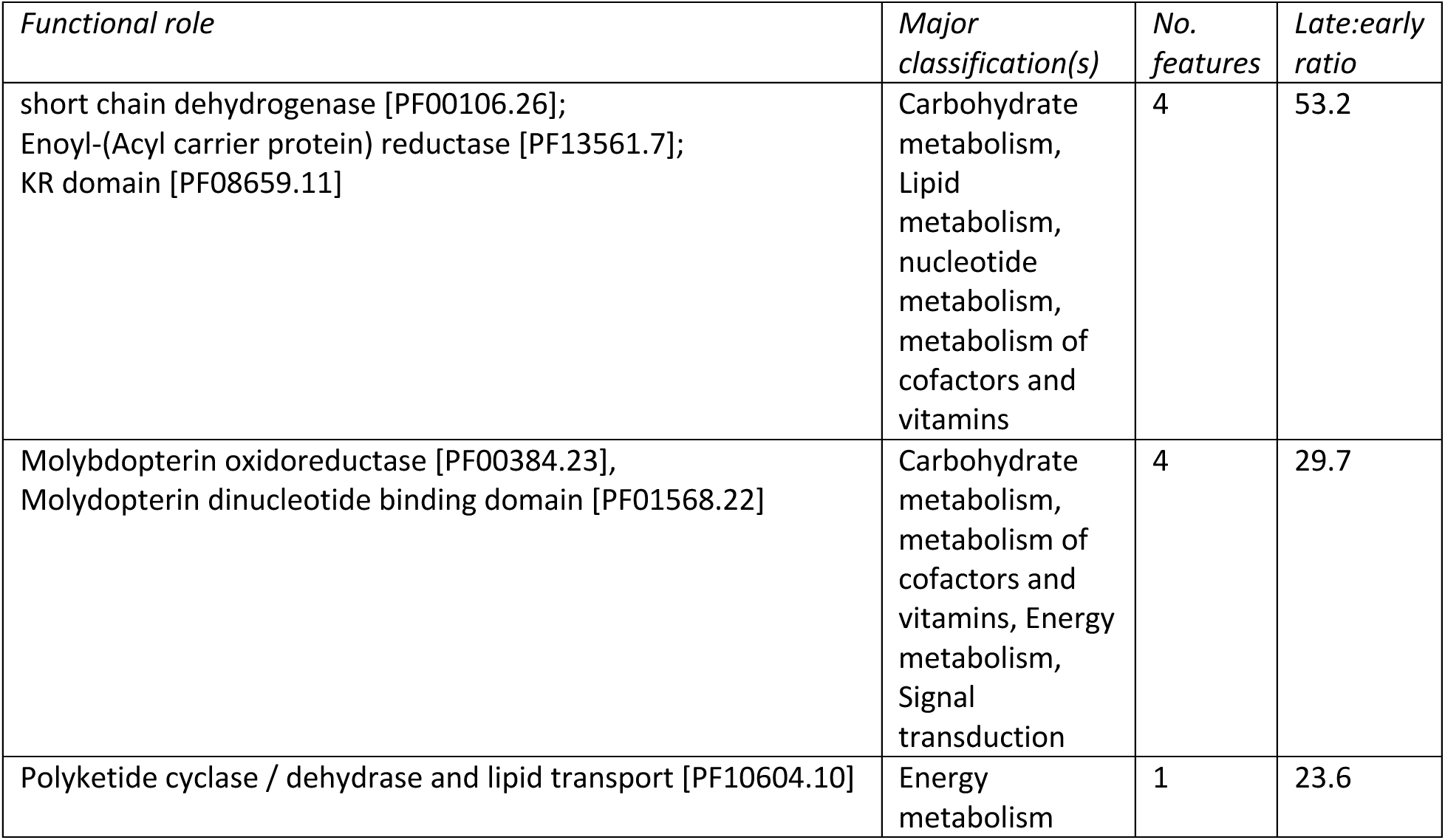
Functional roles that exhibited strong seasonality with transcript enrichment by more than 20-fold in the late season than early. These roles were consistently detected among phyllosphere focal MAGs (39/40 detections).

| *Functional role* | *Major classification(s)* | *No. features* | *Late:early ratio* |
| --- | --- | --- | --- |
| short chain dehydrogenase [PF00106.26];<br>Enoyl-(Acyl carrier protein) reductase [PF13561.7];<br>KR domain [PF08659.11] | Carbohydrate metabolism,<br>Lipid metabolism,<br>nucleotide metabolism,<br>metabolism of cofactors and vitamins | 4 | 53.2 |
| Molybdopterin oxidoreductase [PF00384.23],<br>Molybdopterin dinucleotide binding domain [PF01568.22] | Carbohydrate metabolism,<br>metabolism of cofactors and vitamins, Energy metabolism,<br>Signal transduction | 4 | 29.7 |
| Polyketide cyclase / dehydrase and lipid transport [PF10604.10] | Energy metabolism | 1 | 23.6 |

We then performed metabolic, biosynthetic, and plant-associated gene pathway analyses to understand the functions detected among focal MAGs in the phyllosphere and their activities inferred by transcript recruitment to the pathways (**Figure 6**). Pathways for terpene metabolism (34/40), betaine biosynthesis (30/40), trehalose metabolism (25/40), cyanide degradation (21/40), ROS degradation (23/40), and indole acetic acid (IAA) degradation (40/40) were the most common pathways shared and active across MAGs, and were found in lineages from all four classes represented by the focal MAGs. Thus, these pathways likely are common among phyllosphere members and highly supportive of a leaf-associated lifestyle. In addition, there were some functional pathways and activities that may be more specialized functions because they were detected in only a few MAGs (e.g., xylitol metabolism). However, there was no clear phylogenetic pattern to the distributions of these putative specialized functions.

**Figure 6.**
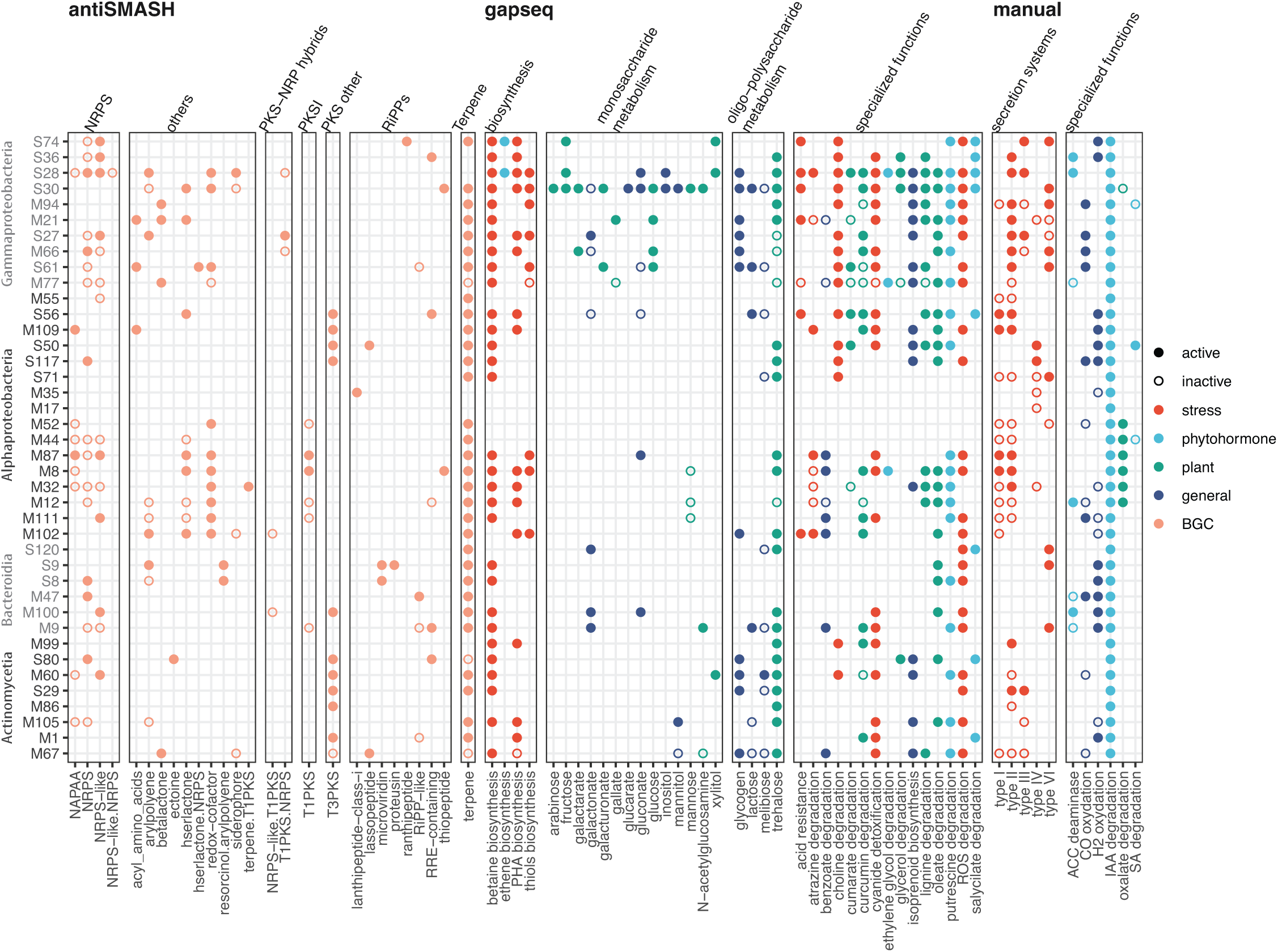
Key functional gene pathways detected in the 40 focal MAGs and their activities (mapped transcripts) during the 2016-2017 switchgrass growing season. Functional gene pathways were curated using antiSMASH for biosynthetic gene clusters, gapseq for general metabolic pathways, and manual selection for plant-associative functions reported in the literature. Pathways that were discovered in the MAGs but not detected in the transcripts are represented by open circles, and pathways detected in the MAGs and mapped by transcripts are represented by filled circles. Colors categorize different functional groups of pathways.

Because of the near-ubiquitous detection of terpene-related biosynthetic genes on the focal MAGs, we examined their annotations and recruitment more closely using gene function predictions (i.e., ORFs), which was expected to improve sensitivity of functional annotation relative to metabolic pathway tools (e.g., BGC). We detected 278 ORFs associated with terpenes annotations in the metatranscripts. Collectively, these terpene-related genes followed the trend of gradual enrichment over the season (**Figure 7A**) and were relatively abundant in both years. The highest enrichment subcategory was terpenoid backbone biosynthesis, which included 29 ORFs. Among several isoprene biosynthesis genes detected, of interest was relatively high, seasonal enrichment of the two terminal enzymes in the non-mevalonate isoprene biosynthesis pathway that is employed by bacteria, the *gcpE* and *lytB* (**Figure 7B, Figure S5**). *GcpE* and *lytB* gene transcripts were detected in more than a third of the MAGs (13/40), and these included MAGs representing all three phyla detected. Six of the eight Proteobacteria that recruited isoprene biosynthesis transcripts were annotated to *Methylobacterium* (alpha). The four Actinomycetia were more distributed phylogenetically and annotated as genera *Frigoribacterium* M1, *Microbacterium* M105, *Amnibacterium* M67, and Pseudokineococcus M86. The single Bacteroidota that had isoprene biosynthesis transcripts was a *Hymenobacter* M9. We then investigated the 40 MAGs for detection of any of the nine genes previously reported to be involved in bacterial biosynthesis of isopentenyl diphosphate, a precursor terpenoids like isoprene (Julsing *et al*., 2007) and found that 29/40 MAGs had six or more of the genes detected, and that all MAGs had at minimum 2 of the genes (**Table S2**). The consistent detection of genes associated with isoprene pathways in these MAGs, which are >50% complete, suggests that biosynthesis of isoprene-related molecules may be a prominent leaf strategy by phyllosphere bacteria. *Psuedomonas* MAG S28, noted previously to be the dominant population that colonized and activated early in the season (**Figure 4, Figure S2**), had high isoprene biosynthesis transcript enrichment early in the season that then declined. However, the other eleven MAGs harboring genes from isoprene biosynthesis pathways compensated with increased activity in the late season.

**Figure 7.**
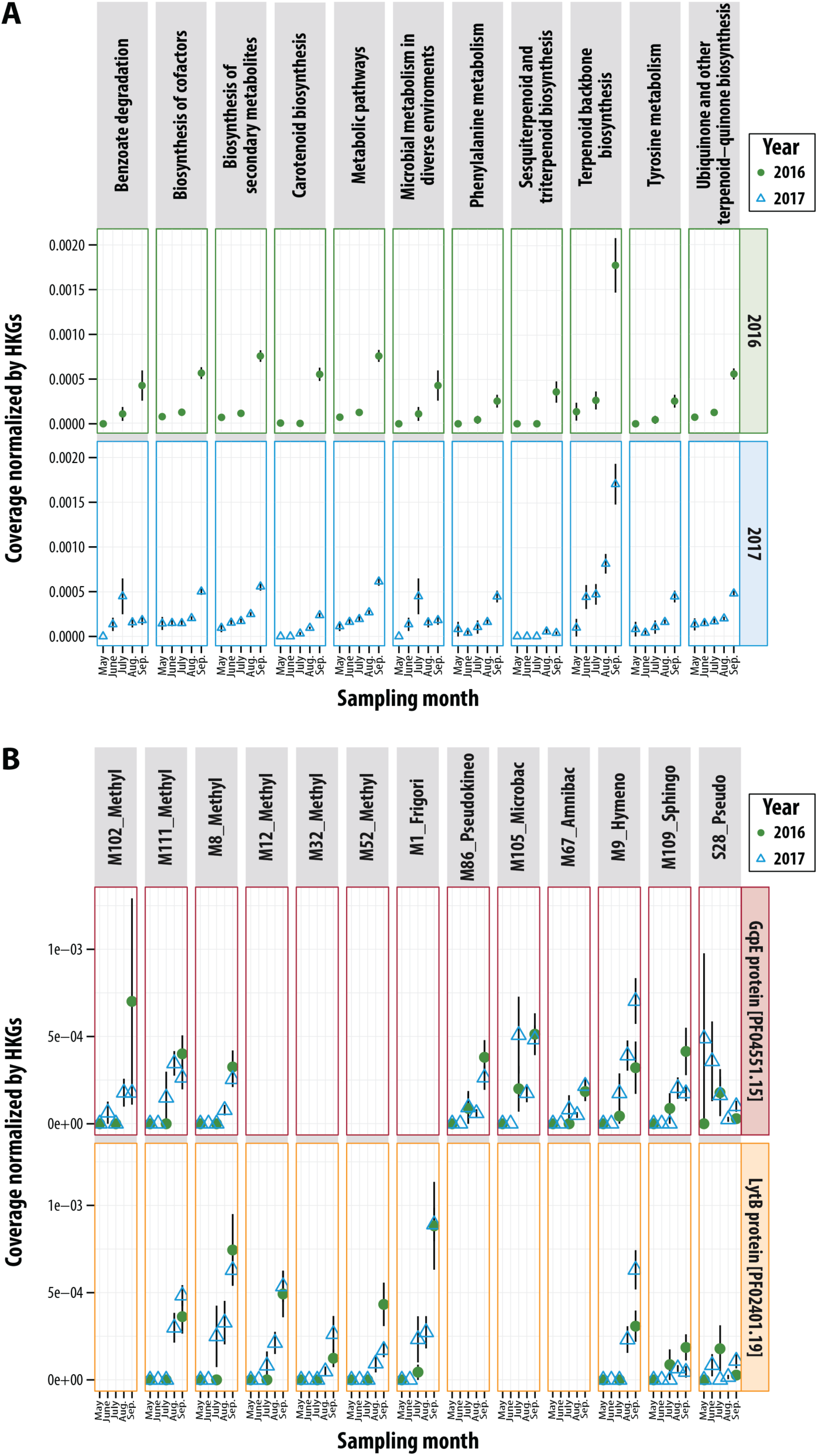
**(A).** 2016 (circle) and 2017 (triangle) switchgrass leaf transcript dynamics of KEGG metabolism classifications associated with terpene metabolism. **(B).** Transcripts in MAGs associated with terminal enzymes in the non-mevalonate isoprene biosynthesis, *gcpE* and *lytB*. MAG IDs include predicted taxonomy at the genus level: Methyl = *Methylobacterium*, Frigor = *Frigoribacterium*, Pseudokineo = *Pseudokineococcus*, Microbac = *Microbacterium*, Amnibac = *Amnibacterium*, Hymeno = *Hymenobacter,* Sphingo = *Sphingomonas,* and Pseudo = *Pseudomonas*.

Returning to the BGC annotations (by antiSMASH, **Figure 6**), most of the terpene transcripts were associated to pigment biosynthesis (e.g., carotenoids). The second most consistent class of BGC transcripts on the focal MAGs were non-ribosomal peptide synthase (NRPK) genes, but the majority of these did not have additional annotations beyond the general category. Therefore, the ORF analysis and BGC detection were complementary in results, especially for terpene-related functions. Another notable BGC finding was that all but two (of eight) Actinomycetia MAGs had transcripts for type III polyketide synthases, while this BGC was less commonly detected among Proteobacteria and Bacteroidota.

### MAGs were detected in metagenomes from different field sites, crops, and years

We next asked if these focal MAGs were detected more broadly in other crop metagenomes. Because leaf metagenomes were not findable, we curated related soil metagenomes from local cropping systems in Michigan and also from the publicly available data within the Integrated Microbial Genomes (IMG) database, which serves as a repository for the Joint Genome Institute’s sequencing efforts through the US Department of Energy. We were surprised to find that, for each metagenome investigated, reads could be mapped to this set of MAGs. This included soil metagenomes from both switchgrass and miscanthus fields, three locations (Iowa, Michigan, and Wisconsin) and sampled across five different years (**Figure 8)**. These results suggest that the detected and analyzed MAGs can be widely distributed in midwestern agroecosystems, and potentially of general importance for perennial crop environments.

**Figure 8.**
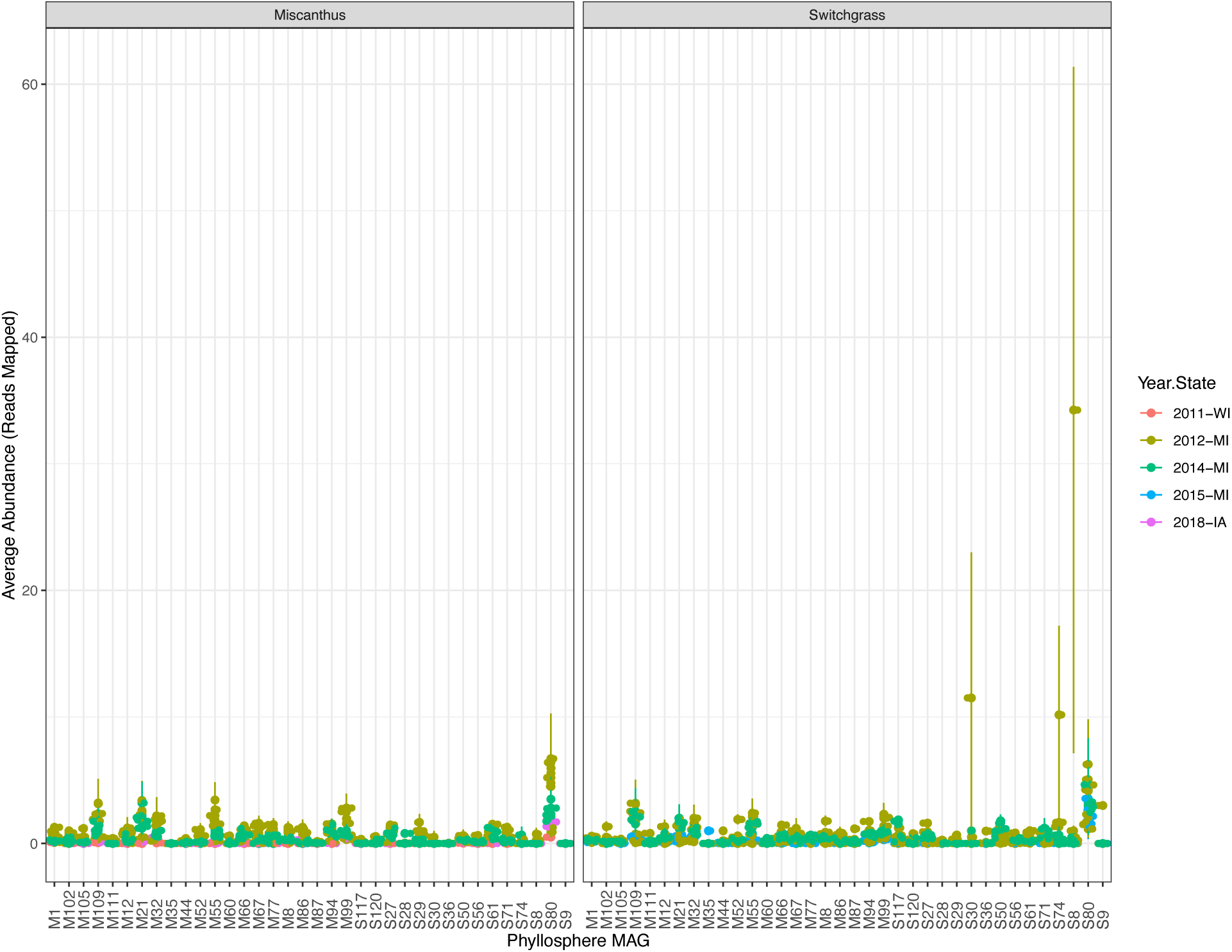
Detection of 36 focal MAGs (out of 40) in publicly available metagenomes and metatranscriptomes of bioenergy grasses.

## Discussion

Here, we report the first multi-year seasonal metagenome and metatranscriptome assessment of plant phyllosphere in agricultural field conditions, focusing on the bacterial functions associated with two promising biofuel feedstocks. We expect these findings to have relevance for other grasses or systems with substantial aerial biomass, including native prairie. Furthermore, our collection of MAGs included phyllosphere members previously reported as abundant, persistent, or key for microbiome assembly in other plants, including the model *Arabidopsis* (e.g., (Carlström *et al*., 2019), specifically: Sphingomonadales, Pseudomonadales, Actinomycetales, Burkholderiales, and Rhizobiales. Therefore, the patterns and consistently detected functions among this curated collection offer general insights as well as seasonal phyllosphere functions.

There are multiple lines of evidence that the focal MAGs discussed here represent key lineages in the switchgrass and miscanthus phyllosphere. First, there is ample overlap among these focal MAGs and the core taxa of high abundance and occupancy from our previous 16S rRNA gene amplicon analysis of the switchgrass and miscanthus phyllosphere diversity (Grady *et al*., 2019), including a MAG associated to *Hymenobacter* M9, which is a genus that had high occupancy across fields and over time, as well as several OTUs assigned. This same time series was investigated in our previous amplicon analysis and the core taxa were prioritized using consistency across replicated fields at the same time point, persistence over time, and relative abundance. This suggests that the populations represented by the MAGs are not rare taxa that are transient to the system. Second, we were able to generate quality assemblies from the complex metagenome data, which is a process that is generally biased towards abundant members. These MAGs are highly represented in read abundance in metagenomes and recruited relatively more metatranscriptome reads as well, suggesting that they are both abundant and active in the phyllosphere. Though it is possible that there are key lineages missing among our collection, we are confident that those discussed here are among the major host- or environment-selected populations inhabiting the switchgrass and miscanthus phyllosphere.

### Stress responses: trehalose, betaine, reactive oxygen, IAA

Trehalose is a disaccharide that protects cells against salt, water, and osmolyte stress by serving as a stabilizing chemical chaperone, either by displacing water from protein surfaces or by vitrifying around protein structures to shield them (Laskowska *et al*., 2020). Similarly, betaine is another commonly biosynthesized osmoprotectant used by microorganisms to contend with water, salt, and temperature stress (Zou *et al*., 2016). Both have been hypothesized to be important survival strategies of microorganisms in the phyllosphere, with supporting evidence from isolate genome analyses (Rastogi *et al*., 2013). Here, we show both trehalose and betaine to be prominent among MAG populations and activated consistently on the leaf surface, suggesting that they are not seasonally activated but rather necessary for the leaf-associated lifestyle.

In bacteria, trehalose metabolism prevents cellular overflow metabolism and carbon stress by redirecting glucose-6-phosphate from conversion to pyruvate (Laskowska *et al*., 2020). Trehalose biosynthesis is common among bacteria and archaea that live in arid, saline, thermal, or seasonally dry environments (e.g., (Urrejola *et al*., 2019), (Dai *et al*., 2015; Lacerda-Júnior *et al*., 2019), and it has also been reported to be induced in a pseudomonad by ethanol (Harty *et al*., 2019), which can originate inside plant cells (Kimmerer and MacDonald, 1987) or more generally in the roots (e.g., Ferner *et al*., 2012), especially during stress, fruit ripening, or senescence (Kimmerer and Kozlowski, 1982). In switchgrass and plants in general, trehalose concentration is increased in response to drought conditions (Liu *et al*., 2015), and its precursor, trehalose-6-phosphate, induces senescence when carbon is readily available (Wingler *et al*., 2012). Furthermore, the *A. thaliana* phyllosphere member *Sphingomonas melonis* was reported to regulate trehalose biosynthesis during growth conditions that promoted mild stress (Gottschlich *et al*., 2019). Given this, it makes sense that the majority of MAGs had enrichment of transcripts related to trehalose metabolism, which would support a plant-associated lifestyle during drought and host senescence (Gottschlich *et al*., 2019).

Betaine biosynthesis in bacteria often begins with oxidation of choline, which is a part of plant tissues and can be transported into the cell (Chen *et al*., 2013). Indeed, choline degradation was detected in 9/10 Gammaproteobacteria MAGs here (**Figure 6**). Microbial-derived osmolytes such as betaine and trehalose have been suggested as targets for biotechnology development to support crop stress tolerance. However, plants can biosynthesize betaine and can be divided into groups of those that do and do not accumulate it in concentrations that are supportive for stress tolerance (Valenzuela-Soto and Figueroa-Soto, 2019). For example, in switchgrass, the concentration of glycine betaine was not predictive of differences in drought tolerance among different genotypes, while trehalose was, along with abscisic acid, spermine, and fructose (Liu *et al*., 2015). Furthermore, our data suggest no notable microbial limitations in the genetic potential or activation of betaine biosynthesis in phyllosphere.

Reactive oxygen species (ROS) serve as signals for various developmental and cellular processes in plants, and here we detected active pathways for ROS degradation in the majority of focal MAGs. Though the precise mechanisms are unclear, homeostasis of ROS are expected to be involved in senescence (Kerchev *et al*., 2015), which is relevant to our study given that at least some to a majority of senescing plants were observed per plot in August and September sampling dates, respectively. Additionally, ROS accumulate in plants that are exposed to abiotic stress, to negative effects. ROS degradation is one of many functions phyllosphere microorganisms employ to contend with expected fluctuations in ROS on the leaf surface (though, these fluxes are difficult to measure (Kerchev *et al*., 2015)). Previously, several genes relevant for oxidative stress response were differentially regulated in a wild-type phyllosphere bacteria *Sphingomonas melonis* strain Fr1 as compared to a knock-out mutant for regulation of general stress response, when both were grown in a medium expected to induce low-levels of stress. (Gottschlich *et al*., 2019). Given that managing plant ROS is a target for reducing crop stress and regulating plant development (Considine and Foyer, 2014; Kerchev *et al*., 2015), it is possible that manipulating microbial ROS degradation could be applied as one tool to achieve such efforts, but much more research is needed to understand the microbial-host interaction given ROS exposure or accumulation, and any possible ROS signaling between them.

IAA is a phytohormone produced by plants to regulate many processes in growth and stress response (e.g., Spaepen *et al*., 2007; Egamberdieva *et al*., 2017). It is also made by many microorganisms, including those shown to support plant grown promotion (e.g., Egamberdieva *et al*., 2017). Therefore, the activity of IAA degradation pathways by focal MAGs is expected given the redundancies between plants and microorganisms in synthesizing and responding to IAA and demonstrates microbiome responsiveness to feedbacks in the host environment.

### Biosynthesis of isoprene-related molecules

Most of the functions identified in our MAGs suggest general requirements for growth and maintenance given a leaf-associated lifestyle (e.g., carbohydrate and amino acid metabolism, pigment production to protect from radiation, similar to previous reports, e.g., Lajoie *et al*., 2020). However, BGC analysis revealed surprising consistency in terpene metabolism pathways, leading us to look more closely at transcript ORFs associated to terpenes. This analysis revealed particular enrichment in pathways and key genes associated with isoprenoid biosynthesis. Isoprenoids are a class of volatile terpenes that are generally abundant and reactive, and they engage in indirect and complex feedbacks with methane and nitrous oxide greenhouse gases (McGenity *et al*., 2018). Isoprene is one of the simplest isoprenoids. It is released by many plant species, and much of it is synthesized within the methylerythritol phosphate pathway of the chloroplast (MEP, aka:non-mevalonate) (Sharkey *et al*., 2008). Isoprene is thought to act as a signaling molecule in stress response (Zuo *et al*., 2019). Studies have also found that isoprene emission protects leaf photosynthesis against short episodes of high-temperature (Sharkey and Yeh, 2001). Plants emit isoprene from matured, photosynthetically active leaves, and emissions are light responsive (Sharkey *et al*., 2008). However, senescing leaves have been reported to decrease in their isoprene emissions relative to leaves at peak growth (Sharkey *et al*., 1991). Both switchgrass and miscanthus have been reported to emit relatively low basal levels of isoprene (Eller *et al*., 2011; Morrison *et al*., 2016).

We hypothesize that that members of the biofuel feedstock phyllosphere bacterial community may be either compensating for the loss of plant-derived isoprene, engaging in interspecies isoprenoid signaling with the host, protecting plant photosynthesis from thermal damage, quenching reactive oxygen species, or possibly producing isoprenoids as overflow metabolites (as hypothesized for *Bacillus subtilus* (Sivy *et al*., 2002)). Bacterial isoprene degraders and synthesizers are widespread in nature (McGenity *et al*., 2018) and have been previously investigated in phyllosphere communities of the relatively high isoprene emitter *Populus* spp (Crombie *et al*., 2018), as well as in soils (El Khawand *et al*., 2016), which can serve as an isoprene sink. Stable isotope assays have been used to determine that a subset of bacteria community members degrade isoprene, including several Actinobacteria (*Rhodococcus* spp.) and *Variovorax* (Proteobacteria) (El Khawand *et al*., 2016; Crombie *et al*., 2018). Our MAG collection contains several Actinomycetia, a *Hymenobacter*, several *Methylobacterium*, and *Pseudomonas* MAG S28 that show activation of genes involved in isoprenoid biosynthesis, and adding support for their involvement in related molecular feedbacks in the phyllosphere. Furthermore, given that a third of our modest collection had activity of key isoprenoid biosynthesis genes, there may be many more leaf bacterial contributors that we did not detect here. In addition, these activities were commonly to three Bacteria phyla, it suggests that biosynthesis of isoprene-related molecules may be a very common phyllosphere microbiome function. As isoprene is a precursor to sidechains needed for several quinones (Nowicka and Kruk, 2010), it could be speculated that leaf bacteria scavange isoprene emitted by the host plant to supplement bacterial synthesis of these sidechains, but then compensate with de novo biosynthesis if host decreases production. We observe that isoprenoid synthesis increases seasonally in the majority of MAGs containing these pathways, and concurrently with when plant isoprene emissions also is expected to decrease, directs future work to understand these dynamics and potential isoprenoid-mediated bacterial-host engagement.

### MAGs of interest

We highlight three MAG populations that were of interest because of their taxonomy, functions, dynamics, or detection. All were shared with taxa in our prior 16S rRNA survey, supporting their inclusion as part of the “core” set that was selected by abundance and occupancy. First, high-quality MAG S28 (>97% complete, <2% contamination) was a prominent pioneer and active colonizer of the leaf (Figure 4 Group 1). MAG S28 is related to *Pseudomonas cerasi*, a species reported to have phytopathogen relatives (Kałużna *et al*., 2016), but we did not note any disease symptoms on the leaves analyzed. This population had expected traits of a strong surface colonizer, including colonization, adaptation and motility subsystems. It also had six pathways related to phytohormone responses (out of 7 total phytohormone pathways observed in these data), including activated ethene biosynthesis, ACC deaminase, and degradation of ethylene glycol, putrescine, salicylate and IAA. These data suggest that S28 has several mechanisms to engage or respond to the host via phytohormones.

Next, MAG M9, identified as *Hymenobacter*, was of interest because it was associated to the most numerous taxonomic group detected in our prior 16S rRNA gene survey (Grady *et al*., 2019) and not among the most typically investigated phyllosphere lineages in the literature. While M9 populations were first detected early in the season, its transcripts were enriched in the late season (**Figure 4** Group 5). MAG M9 had detected and activated galactonate, N-acetylglucosamime, and lactose metabolisms, which were not common among the focal MAGs. It also had activated benzoate, curcumin, and putrescine degradation, as well as cyanide detoxification, type VI secretion, and dihydrogen oxidation. While M9 also had some pathways that were common among these MAGs (e.g., ROS and IAA degredation, terpene biosynthesis), its suite of more sparsely detected pathways and functions suggest a specialized role in the phyllosphere community. Notably, M9 has 65% completion and 0% detected contamination, suggesting more functional potential remains to be discovered for this and similar *Hymenobacter* lineages inhabiting the phyllosphere.

Finally, we selected a representative Actinomycetia MAG M60, a *Quadrisphaera* lineage that had activated isoprenoid biosynthesis and had increased activity late in the season along with the majority of focal MAGs (Figure 4, Group 3). Studies have found that members of Actinomycetia are important part of phyllosphere that contribute to disease prevention and plant growth [El-Tarabily 2009; Javed 2021; Anwar 2016]. MAG M60 had several oligo/polysaccharide metabolisms that were infrequently detected in these data, including glycogen, melibiose, and trehalose. Despite its high completeness and low contamination (>95% and < 5%, respectively), M60 was sparsely annotated by the methods we applied. However, *Quadrisphaera* have been reported to be highly abundant in the phyllosphere or endosphere of various plants (Bao *et al*., 2020).

### Conclusions

Many recent review, perspective, and opinion pieces have urged integration of multi-omics approaches to improve understanding of the microbiome and its relationship to the host plant (Rastogi *et al*., 2013; Levy *et al*., 2018; Remus-Emsermann and Schlechter, 2018; Beilsmith *et al*., 2019; Trivedi *et al*., 2020). However, most integrative studies have focused almost exclusively on the rhizosphere as the compartment of soil-plant feedbacks and nutrient and water acquisition for the host. Though leaves are readily accessible for sampling, the phyllosphere microbiome is challenging to investigate using throughput, cultivation-independent approaches like metagenomics and metatranscriptomics. There are high levels of host and chloroplast contamination in leaf samples, and relatively low microbial biomass per leaf that that must be first dislodged from tightly-adhered biofilms. Signal from messenger RNA in metatranscriptome analysis is masked by abundant ribosomal RNA signal, leading to further challenge. Because of the combination of all of these challenges, much of our understanding of the phyllosphere, as the largest surface area of microbial habitation on Earth (Peñuelas and Terradas, 2014), has been learned from studies that employ model hosts and synthetic or model microbial communities in controlled settings, or from description of the community structure by sequencing of marker genes, amplified and bioinformatically depleted of chloroplast genes to overcome the challenges of low signal and host contamination.

Here, we report optimized laboratory protocols (to minimize host and chloroplast signals) combined with a genome-centric bioinformatic approach to perform focused functional gene and transcript analysis of seasonally dynamic yet persistent phyllosphere microbiome members. To our knowledge, this is the first untargeted bacterial metatranscriptomic work performed on the leaf phyllosphere of field-grown crops. Other recent leaf metatranscriptome studies have focused the viral communities of tomato and pepper (Choi *et al*., 2020), soybean (Marzano and Domier, 2016), and rice (Chao *et al*., 2020 Preprint). A key strength of this work is the challenging integration of phyllosphere metagenome and metatranscriptome data, leveraging the higher coverage of the metagenomes with the activity information available from the metatranscriptomes. Despite the relatively limited coverage of the MAGs (due to substantial host and ribosomal DNA contamination), the analysis proved successful by integrating both datasets and focusing on genome-centric interpretation. Thus, there are likely many more prevalent and functionally active populations of the phyllosphere that were not captured in this study, including those players previously known to be key in the phyllosphere (e.g. Delmotte *et al*., 2009; Vorholt, 2012). Substantial additional sequencing effort or an enrichment strategy would improve signal for a cultivation-independent approach to target those players. While the use of genome-centric approaches has the obvious shortcoming that we have obviously not captured every microbiome member, our approach does allow us to link actively transcribed functions to specific microbial membership. Furthermore, the functional genes and activities documented here are logical given current understanding of microbial adaptation to the host and phyllosphere environment.

Overall, this work provides evidence of a thriving, dynamic, functionally diverse, leaf-specialized, and host-responsive microbiome on the phyllosphere of perennial grasses. It provides evidence of specific phyllosphere functions that are seasonally activated in a temperate agroecosystem and suggests several hypotheses of important host-microbe interactions in the phyllosphere, for example via central metabolism, isoprenoid biosynthesis, and stress response engagements. This research contributes to our broad understanding of the dynamics and activities of phyllosphere microbial communities, and points to specific microbial functions to target that could prove useful for plant-microbiome management.

## Acknowledgements

Support for this research was provided by the Great Lakes Bioenergy Research Center, U.S. Department of Energy, Office of Science, Office of Biological and Environmental Research (Awards DE-SC0018409 and DE-FC02-07ER64494), by the National Science Foundation Long-term Ecological Research Program (DEB 1637653 and 1832042) at the Kellogg Biological Station, Michigan State University AgBioResearch, and by the DOE Center for Advanced Bioenergy and Bioproducts Innovation (US Department of Energy, Office of Science, Office of Biological and Environmental Research under award number DE-SC0018420). This work was also supported in part by Michigan State University through computational resources provided by the Institute for Cyber-Enabled Research and in part by the University of Wisconsin-Madison Wisconsin Energy Institute as supported by GLBRC Information Services. The work (proposal:10.46936/10.25585/60000818) conducted by the U.S. Department of Energy Joint Genome Institute (https://ror.org/04xm1d337), a DOE Office of Science User Facility, is supported by the Office of Science of the U.S. Department of Energy operated under Contract No. DE-AC02-05CH11231. AS acknowledges support by the USDA National Institute of Food and Agriculture and Michigan State University AgBioResearch. NS acknowledges support from the Michigan State University Plant Resilience Institute. We thank three anonymous reviewers for their thoughtful comments on a previous version of this work.

## Author contributions

KLG and AS conceived and designed experiments; NS, KLG and AS performed the experiments; ACH, NS, SKD, FY and AS analyzed the data; ACH, NS, SKD, FY, KLG and AS contributed materials/analysis tools; and ACH, NS, SKD, FY, KLG and AS wrote the paper.

## Competing Interests statement

The authors declare no competing interests.

## Datasets

**Dataset 1.** Metadata for metagenome samples according to MIMs standards.

**Dataset 2.** Metadata for metatranscriptome samples according to MIMs standards.

**Dataset 3.** Excel file. *Sheet 1.* Metadata for selected focal 40 metagenome-assembled genomes (>50% complete, <10% contamination) according to MiMAG standards *Sheet 2.* Other MAGs (non-focal) assembled from switchgrass or miscanthus metagenomes.

**Dataset 4**. Excel file. *Sheet 1.* Summary of metagenome reads mapped to the focal MAGs. *Sheet 2.* Summary of metatranscriptome reads mapped to the focal MAGs. *Sheet 3*. Total reads mapped from each metagenome. *Sheet 4.* Total reads mapped from each metatranscriptome. **Dataset 5.** Public metagenomes used for comparison to the switchgrass and miscanthus focal MAGs.

**Dataset 6.** Core KEGG functions identified in at least 39 focal MAGs and their estimated abundances in early and late season.

## Supplementary Figures

**Figure S1.** Contaminating plant or fungal sequences. Proportion of sequencing in metagenomes originating from miscanthus and switchgrass phyllospheres associated with miscanthus and switchgrass host genomes (A) and prevalent fungal genomes (B, Table S1). Proportion of sequencing in metatreanscriptomes originating from and switchgrass phyllospheres associated with switchgrass host genomes (C) and fungal genomes (D, Table S1).

**Figure S2.** Hierarchical clustering identified five clusters of MAGS to identify MAG populations with coherent seasonal activity dynamics. Clustering was based on metatranscriptome diversity and abundances.

**Figure S3.** Mean transcript dynamics by month and year for each MAG cluster. Filled circles indicate that identification of transcripts, and empty circles indicate no detection.

**Figure S4.** Summary of transcript seasonality of phyllosphere open reading frames (ORFs) that could be annotated as KEGG functional roles and were consistently detected among focal MAGs (at least 39/40 detections). Ratios are late-to-early normalized transcript abundances on MAGs.

**Figure S5.** 2016 (circle) and 2017 (triangle) switchgrass leaf transcript dynamics of KEGG metabolism classifications associated with terpenoid backbone biosynthesis.

## Supplementary Tables

**Table S1.**
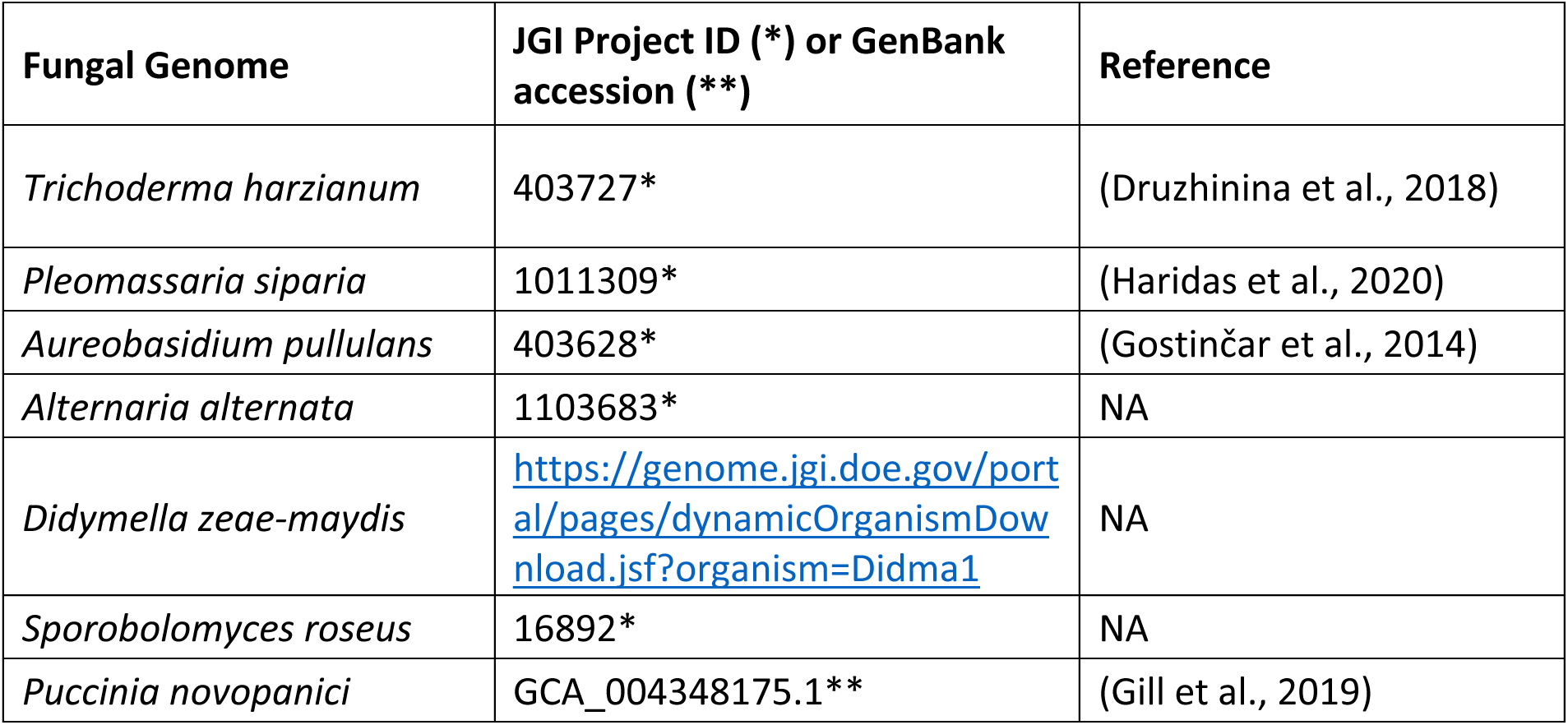
Fungal genomes used for filtering metagenome reads to remove eukaryotic contamination. Genomes were selected to use for filtering based on the taxonomic identities of prevalent fungal taxa detected in our previous ITS2 amplicon survey that was conducted at the same location (Bowsher *et al*., 2020).

**Table S2.**
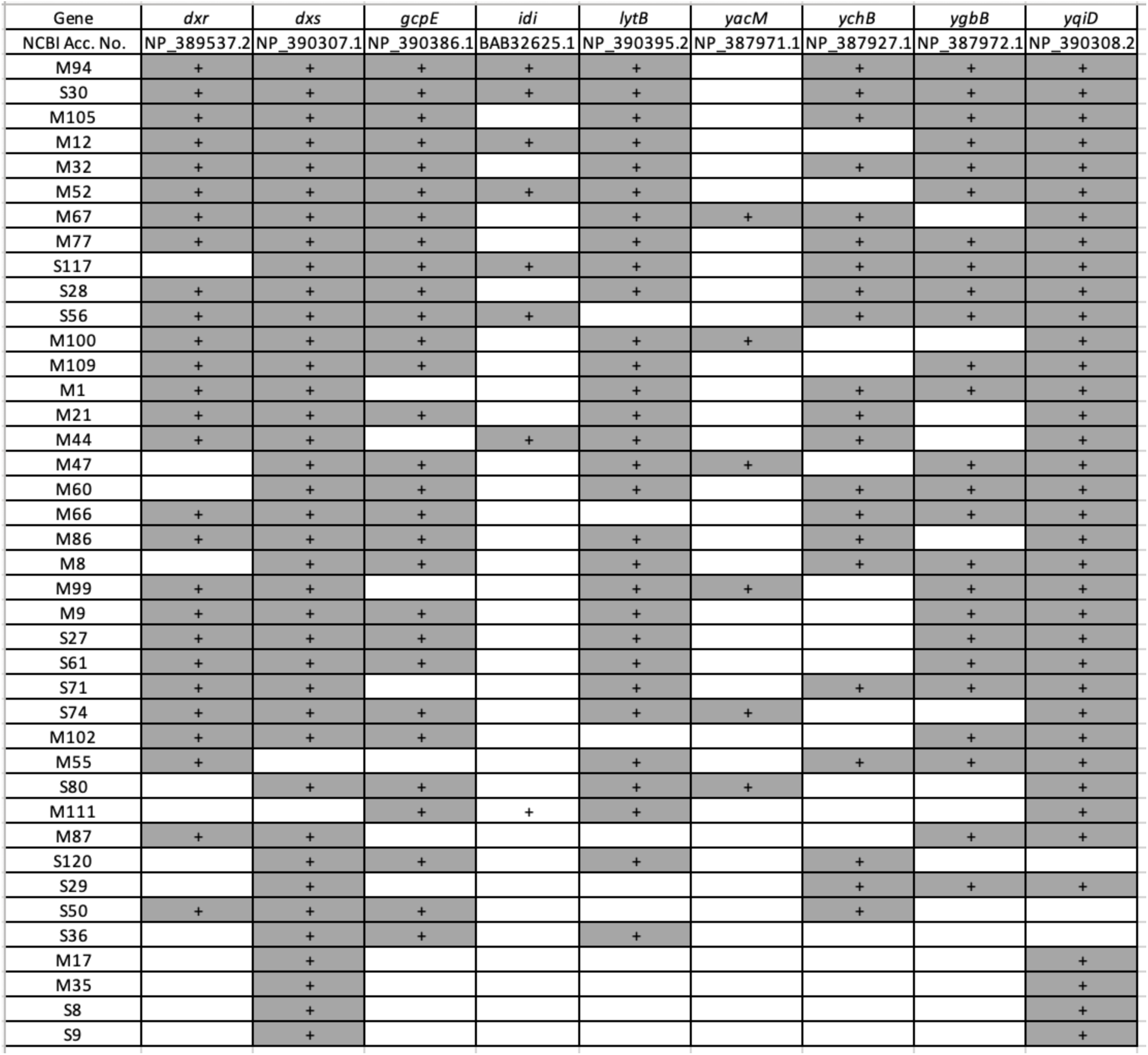
Summary of the nine genes involved the *Bacillus subtilus* isoprene biosynthesis pathway that were detected among focal MAG contigs. Except for *ypgA*, these genes were directly linked to isoprene accumulation by (Julsing *et al*., 2007).

## Notes

https://genome.jgi.doe.gov/portal/Seadynanfunction/Seadynanfunction.info.html

https://github.com/ShadeLab/PAPER_Howe_2021_switchgrass_MetaT

### Competing Interest Statement

The authors have declared no competing interest.

### Summary of Updates

This revision was made in response to constructive comments from three anonymous reviewers (thank you!). Overarching temporal patterns and conclusions from the original study were in general robust to these improvements, which we feel strengthens the work. Specifically, this study has been updated and re-analyzed to: 1) include more MAGs than the original version based on 50% complete and <10% contaminated (omitting the previous inclusion criteria for occupancy); 2) clarify the metadata, code, and methods and make supplemental data files more findable; 3) compare MAGs with previous core microbiome study based on 16S to cross-validate detection of key members; 4) improved and more comprehensive annotation methods for functional genes based on KEGG plus multiple tools; and 5) different transcript standardization, now relative to housekeeping genes.

