## Supplemental Figures and Tables for "Life on the leaf: Seasonal activities of the phyllosphere microbiome of perennial crops"

Figure S1.

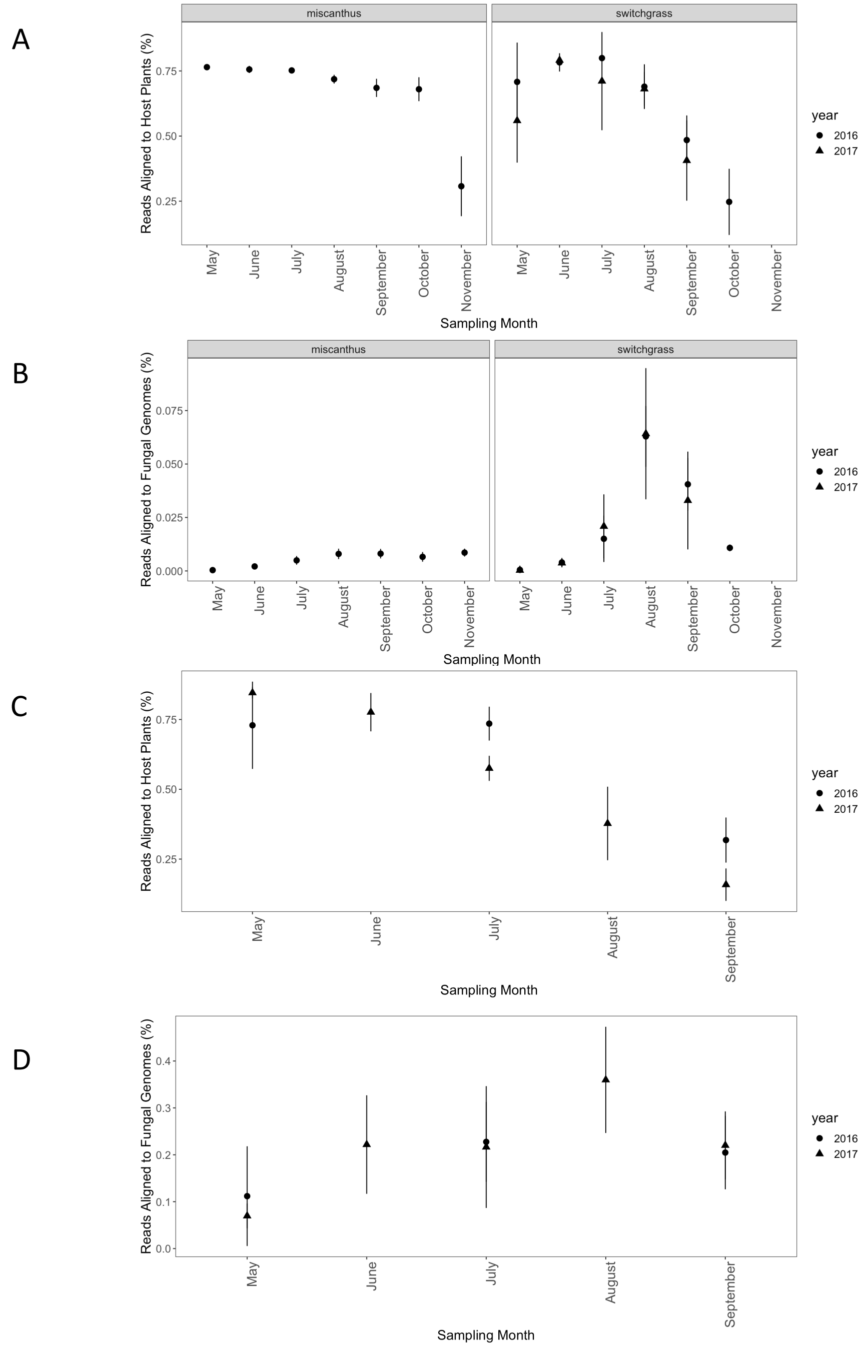


**Figure S1**. Contaminating plant or fungal sequences. Proportion of sequencing in metagenomes originating from miscanthus and switchgrass phyllospheres associated with miscanthus and switchgrass host genomes (A) and prevalent fungal genomes (B, Table S1). Proportion of sequencing in metatreanscriptomes originating from and switchgrass phyllospheres associated with switchgrass host genomes (C) and fungal genomes (D, Table S1).

Figure S2

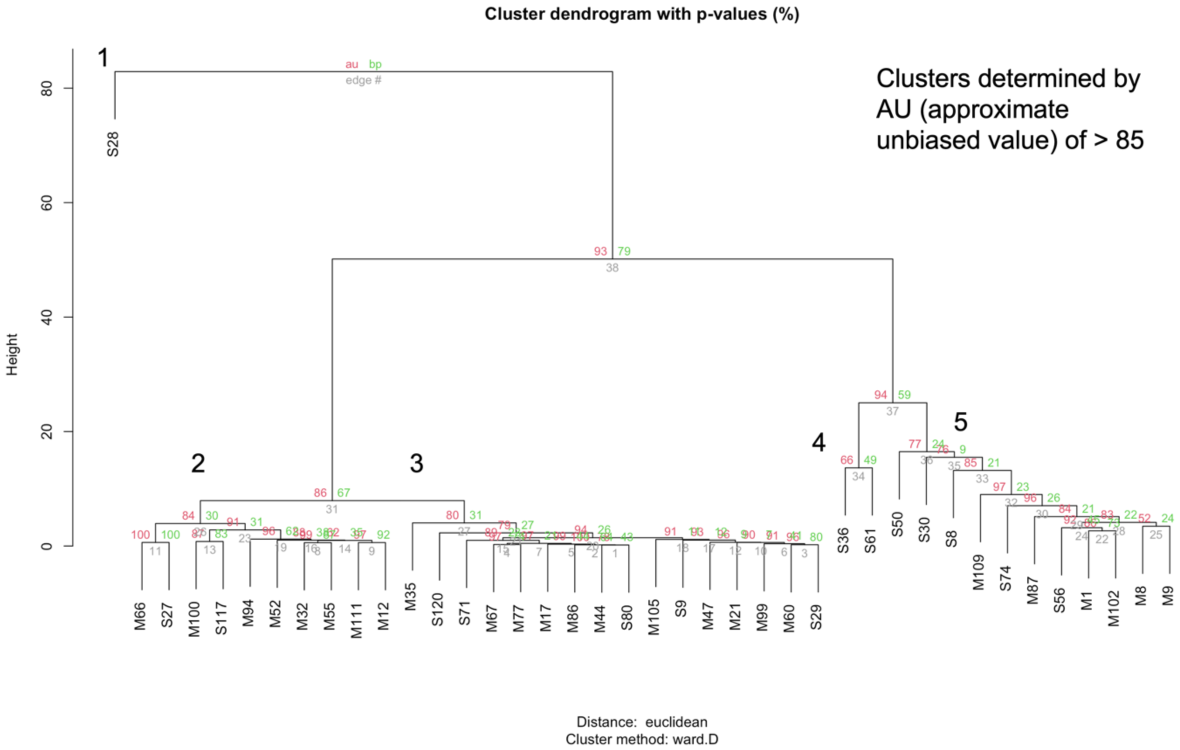


**Figure S2.** Hierarchical clustering identified five clusters of MAGS to identify MAG populations with coherent seasonal activity dynamics. Clustering was based on metatranscriptome diversity and abundances.

Figure S3

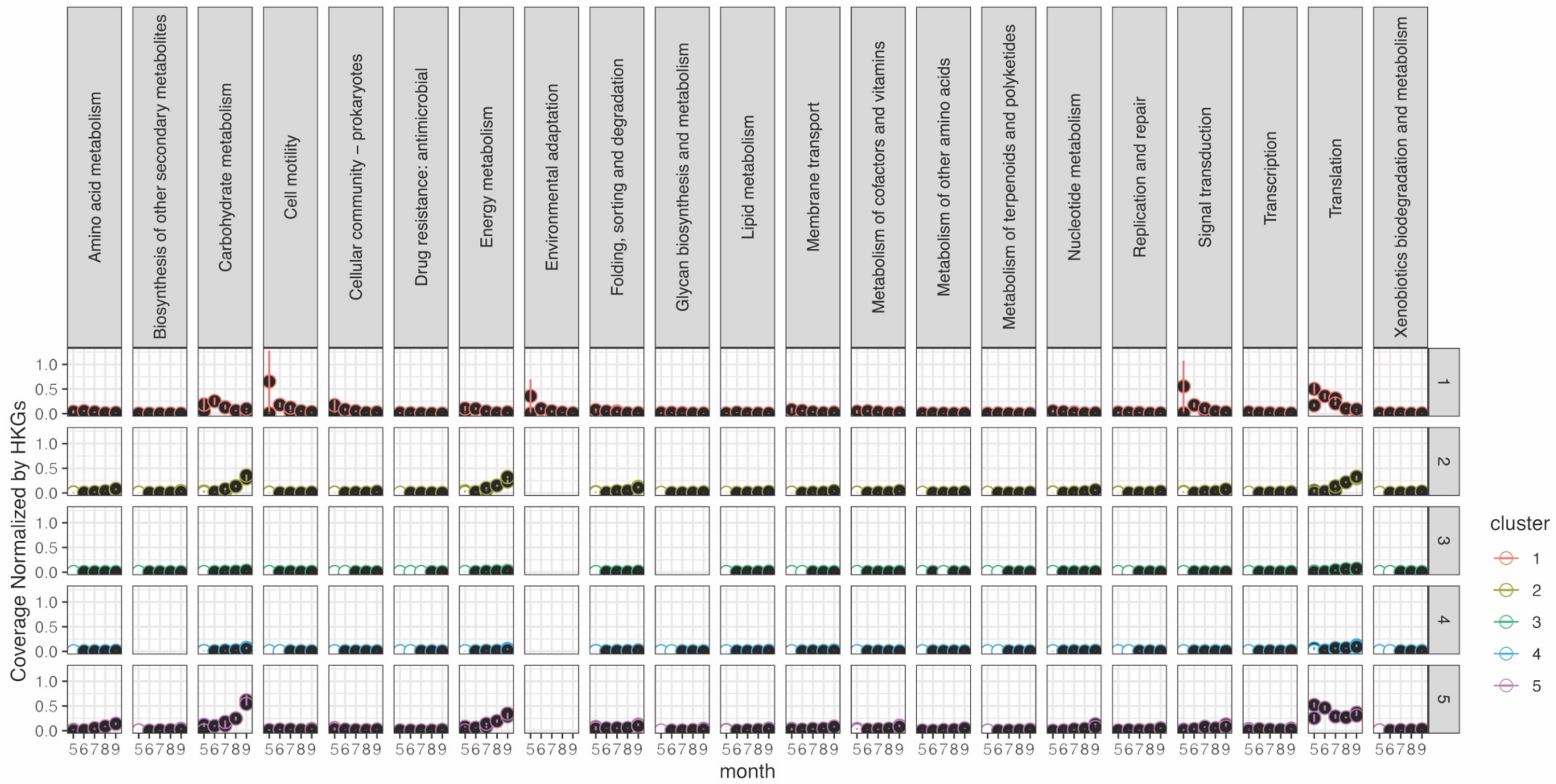


**Figure S3.** Mean transcript dynamics by month and year for each MAG cluster. Filled circles indicate that identification of transcripts, and empty circles indicate no detection.

Figure S4

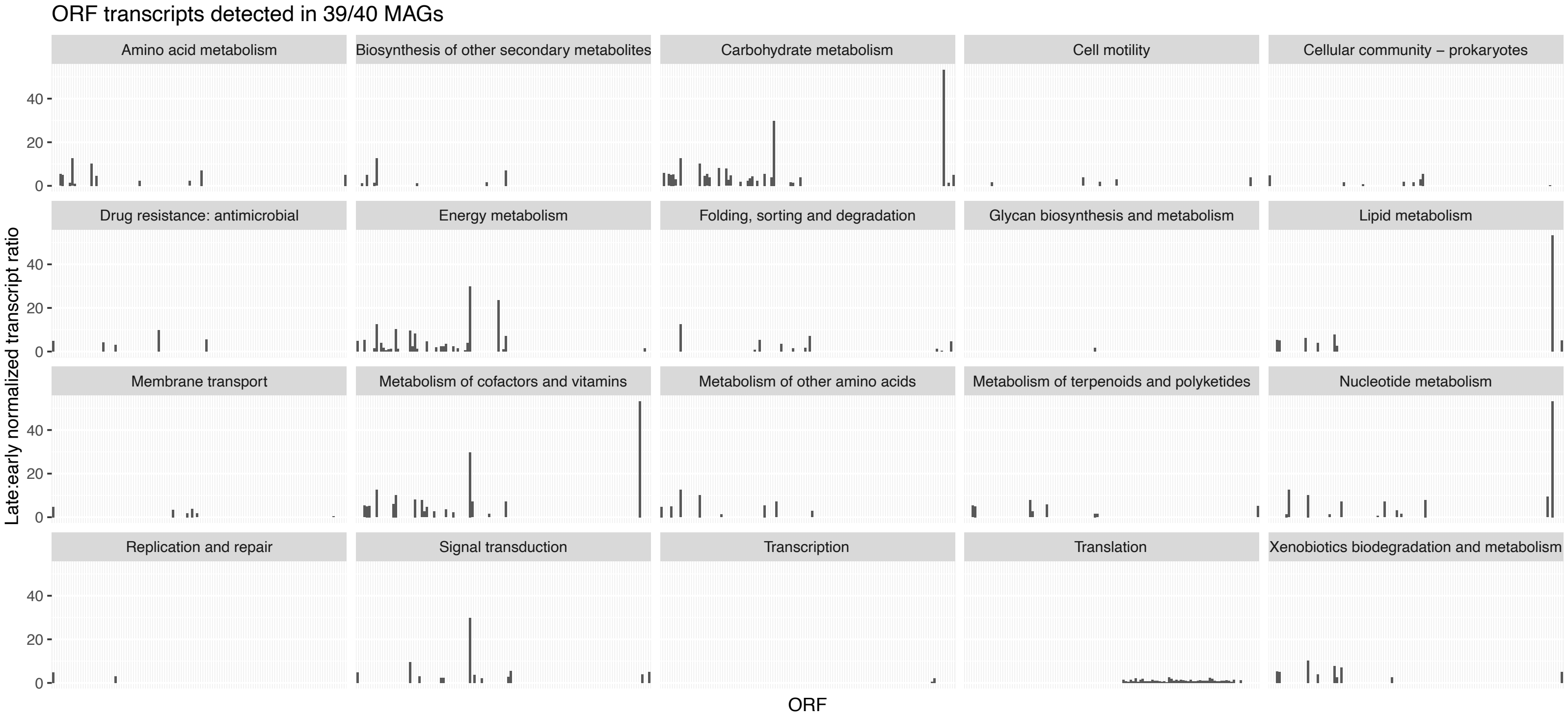


**Figure S4**. Summary of transcript seasonality of phyllosphere open reading frames (ORFs) that could be annotated as KEGG functional roles and were consistently detected among focal MAGs (at least 39/40 detections). Ratios are late-to-early normalized transcript abundances on MAGs.

Figure S5

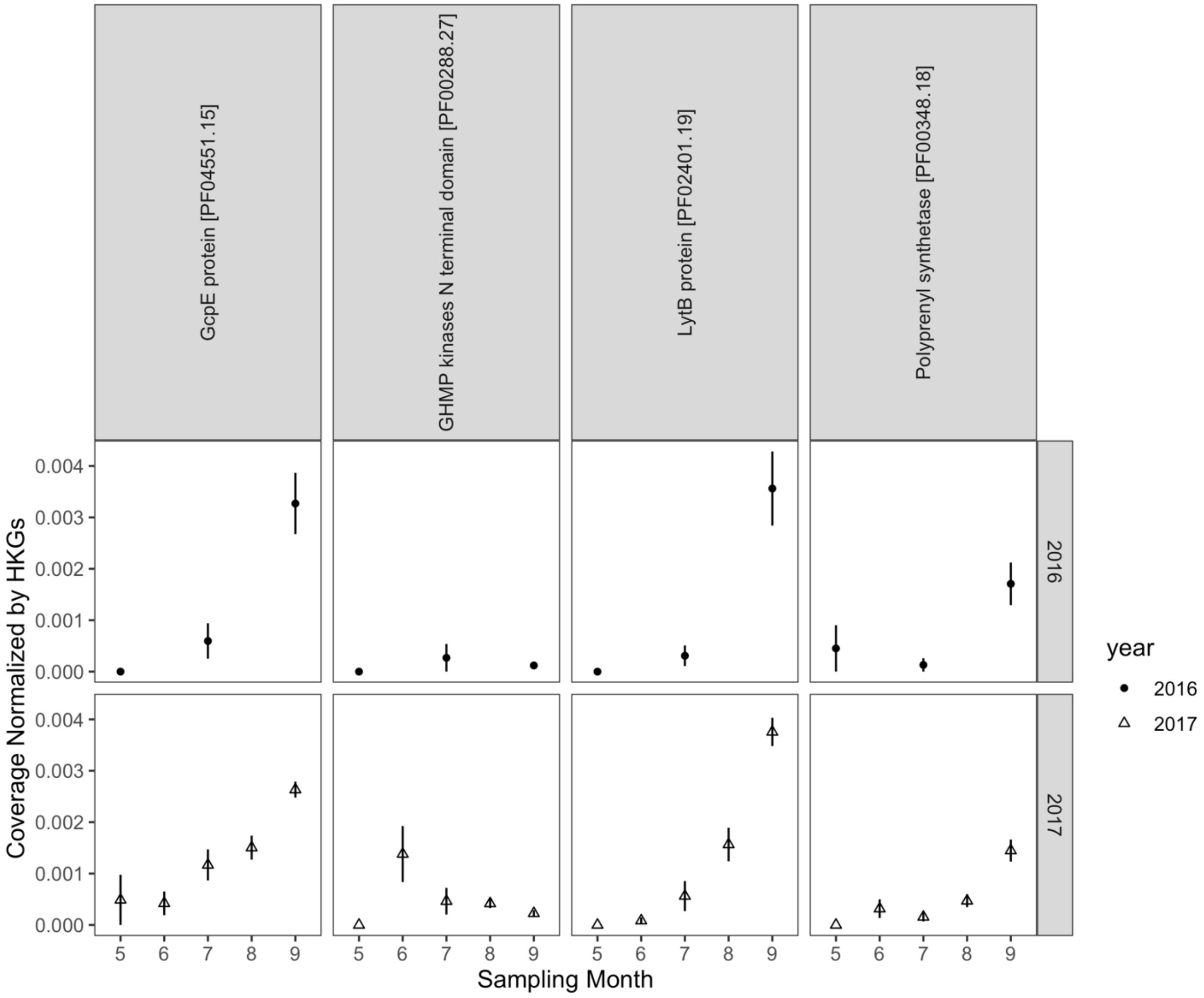


**Figure S5**. 2016 (circles) and 2017 (triangles) switchgrass leaf transcript dynamics of KEGG metabolism classifications associated with terpenoid backbone biosynthesis.

**Table S1**.
Fungal genomes used for filtering metagenome reads to remove eukaryotic contamination. Genomes were selected to use for filtering based on the taxonomic identities of prevalent fungal taxa detected in our previous ITS2 amplicon survey that was conducted at the same location (Bowsher et al., 2020).

| **Fungal Genome** | **JGI Project ID (*) or GenBank accession (**)** | **Reference** |
|---|---|---|
| *Trichoderma harzianum* | 403727* | (Druzhinina et al., 2018) |
| *Pleomassaria siparia* | 1011309* | (Haridas et al., 2020) |
| *Aureobasidium pullulans* | 403628* | (Gostinčar et al., 2014) |
| *Alternaria alternata* | 1103683* | NA |
| *Didymella zeae-maydis* | https://genome.jgi.doe.gov/portal/pages/dynamicOrganismDownload.jsf?organism=Didma1 | NA |
| *Sporobolomyces roseus* | 16892* | NA |
| *Puccinia novopanici* | GCA_004348175.1** | (Gill et al., 2019) |

**Table S2.**

Summary of the nine genes involved the *Bacillus subtilus* isoprene biosynthesis pathway that were detected among focal MAG contigs. Except for *ypgA*, these genes were directly linked to isoprene accumulation by (Julsing *et al.*, 2007).

| Gene | *dxr* | *dxs* | *gcpE* | *idi* | *lytB* | *yacM* | *ychB* | *ygbB* | *yqiD* |
|---|---|---|---|---|---|---|---|---|---|
| NCBI Acc. No. | NP_389537.2 | NP_390307.1 | NP_390386.1 | BAB32625.1 | NP_390395.2 | NP_387971.1 | NP_387927.1 | NP_387972.1 | NP_390308.2 |
| M94 | + | + | + | + | + | | + | + | + |
| S30 | + | + | + | + | + | | + | + | + |
| M105 | + | + | + | | + | | + | + | + |
| M12 | + | + | + | + | + | | | + | + |
| M32 | + | + | + | | + | | + | + | + |
| M52 | + | + | + | + | + | | | + | + |
| M67 | + | + | + | | + | + | + | | + |
| M77 | + | + | + | | + | | + | + | + |
| S117 | | + | + | + | + | | + | + | + |
| S28 | + | + | + | | + | | + | + | + |
| S56 | + | + | + | + | | | + | + | + |
| M100 | + | + | + | | + | + | | | + |
| M109 | + | + | + | | + | | | + | + |
| M1 | + | + | | | + | | + | + | + |
| M21 | + | + | + | | + | | + | | + |
| M44 | + | + | | + | + | | + | | + |
| M47 | | + | + | | + | + | | + | + |
| M60 | | + | + | | + | | + | + | + |
| M66 | + | + | + | | | | + | + | + |
| M86 | + | + | + | | + | | + | | + |
| M8 | | + | + | | + | | + | + | + |
| M99 | + | + | | | + | + | | + | + |
| M9 | + | + | + | | + | | | + | + |
| S27 | + | + | + | | + | | | + | + |
| S61 | + | + | + | | + | | | + | + |
| S71 | + | + | | | + | | + | + | + |
| S74 | + | + | + | | + | + | | | + |
| M102 | + | + | + | | | | | + | + |
| M55 | + | | | | + | | + | + | + |
| S80 | | + | + | | + | + | | | + |
| M111 | | | + | + | + | | | | + |
| M87 | + | + | | | | | | + | + |
| S120 | | + | + | | + | | + | | |
| S29 | | + | | | | | + | + | + |
| S50 | + | + | + | | | | + | | |
| S36 | | + | + | | + | | | | |
| M17 | | + | | | | | | | + |
| M35 | | + | | | | | | | + |
| S8 | | + | | | | | | | + |
| S9 | | + | | | | | | | + |
